# Factors affecting macromolecule orientations in thin films formed in cryo-EM

**DOI:** 10.1101/2023.11.06.565913

**Authors:** Swati Yadav, Kutti R. Vinothkumar

## Abstract

The formation of a vitrified thin film embedded with randomly oriented macromolecules is an essential prerequisite for cryogenic sample electron microscopy. Most commonly, this is achieved using the plunge freeze method described nearly 40 years ago. Although this is a robust method, the behaviour of different macromolecules shows great variation upon freezing and often needs to be optimized to obtain an isotropic, high-resolution reconstruction. For a macromolecule in such a film, the probability of encountering the air-water interface in the time between blotting and freezing and adopting preferred orientations is very high. 3D reconstruction using preferentially oriented particles often leads to anisotropic and uninterpretable maps. Currently, there are no general solutions to this prevalent issue, but several approaches largely focusing on sample preparation with the use of additives and novel grid modifications have been attempted. In this study, the effect of physical and chemical factors on the orientations of macromolecules was investigated through an analysis of select well-studied macromolecules and important parameters that determine the behaviour of proteins on cryo-EM grids were revealed. These insights highlight the nature of interactions that cause preferred orientations and can be utilized to systematically address orientation bias for any given macromolecule and provide a framework to design small-molecule additives to enhance sample stability and behaviour.

## 1. Introduction

Single particle cryo-EM has become an indispensable technique in structural biology owing to its relative ease of image acquisition, reconstruction, and structure determination. Consequently, there has been a steady increase in the number of structures deposited in the Electron Microscopy Database (EMDB) in the past few years (Patwardhan, 2017). Near atomic resolution maps of biological macromolecules can now be routinely obtained due to improved hardware and specimen preparation methods along with developments in algorithms for image processing (Bai, 2021; Y. Cheng, 2015; Chua et al., 2022; Lyumkis, 2019; Nogales, 2015; Vinothkumar & Henderson, 2016; Zheng et al., 2023). Obtaining a thin film of purified macromolecules in a vitreous layer of ice transparent to electrons is an important first step in cryo-EM structure determination. Currently, the plunge freeze method is the most commonly used technique for specimen preparation. This method involves placing a drop of the sample on a plasma-cleaned metal grid (typically coated with carbon and has an array of holes) on which a thin film is formed as the excess fluid is blotted away using blotting paper (Dubochet et al., 1982; Dubochet & McDowall, 1981). The grid is then plunged into a cryogen, causing the vitrification of water and preserving the native structures of macromolecules (Dubochet et al., 1985; Dubochet & McDowall, 1981; Taylor & Glaeser, 1974).

The freezing and blotting conditions must be extensively optimised for temperature and humidity to reproducibly create a single layer of macromolecules on a thin film of ice suitable for imaging. Because of their large size, the first test specimens used to optimise these conditions were liposomes and viruses. They lose their structural integrity if the osmolarity of the buffer changes owing to slow cooling (Adrian et al., 1984; Dubochet et al., 1988). It has been shown through recent studies that during freezing, very often macromolecules can interact with the air-water interface (AWI). This interaction can cause the macromolecules to denature, dissociate, adapt preferential orientations, or avoid the holes completely and stick to the carbon support (D’Imprima et al., 2019; Drulyte et al., 2018; Kampjut et al., 2021; Liu & Wang, 2023; Noble, Dandey, et al., 2018; Noble, Wei, et al., 2018). Along with protein biochemistry, the propensity of macromolecules to adopt preferential orientations is perhaps a common rate-limiting step in high-resolution structure determination using cryo-EM.

Depending on the instrumental setup, the time between blotting and freezing can vary, but it is typically around a second. During this period, the macromolecules are suspended in an extremely thin film and constantly tumble in a solution, undergoing Brownian motion. An intact macromolecule undergoing diffusion can encounter and get trapped in specific orientations at the freshly formed AWI. This effect is protein-dependent and more severe in some cases than in others (Naydenova & Russo, 2017).

When and how does having preferred orientations become a problem? There are cases where the preferred orientation exists but does not pose a problem, such as in proteasomes and viruses, because of their highly symmetric architecture (Campbell et al., 2015; Vogel et al., 1986). A characteristic feature of maps reconstructed from particles with orientation bias is the stretching of density in the cryo-EM map in one direction (anisotropy) (Figure 1), which makes the maps uninterpretable (Sorzano et al., 2022). In such cases, the resolution estimate based on the comparison of the half-maps is unrealistic, and the features in the map do not justify the resolution. Two examples of macromolecules exhibiting orientation bias are shown in Figure 1. It should be noted that 2D projections of different views of a macromolecule contribute differently to the final reconstruction, and this is an important criterion for obtaining a reconstruction with isotropic resolution, as shown by (Tan et al., 2018) and is further demonstrated using catalase as an example in Figure S1.

**Figure 1:**
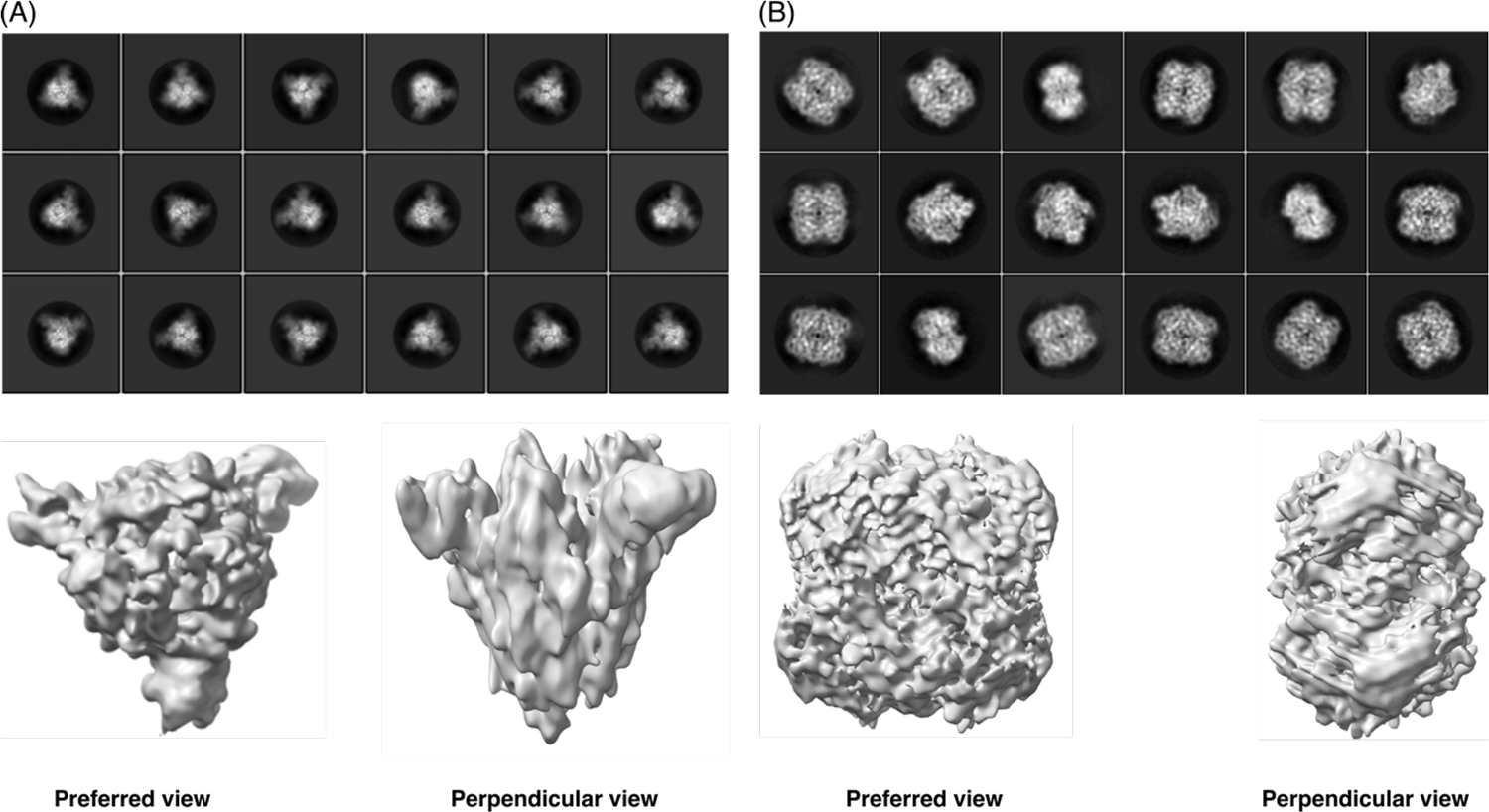
Examples of anisotropic cryo-EM maps resulting from orientation bias. The upper panel shows the reference-free 2D class averages of (A) SARS-CoV-2 spike protein and (B) human erythrocyte catalase. For the spike protein, the preferred bottom views are observed. In the case of catalase, preference for the top/bottom view is evident. In the lower panel, 3D maps with anisotropic features are shown with the preferred and perpendicular views as labelled. The symmetries applied during reconstruction were C1 and D2 for the spike protein and catalase, respectively.

Several approaches have been proposed to address the preferred orientation problem (Drulyte et al., 2018). One such approach is to change the sample preparation process by skipping the blotting with filter papers altogether and minimising the time between thin film creation and freezing to as little as 100 ms. This method uses pL to nL quantities of the sample, which is sprayed onto a self-wicking grid that contains nanowires to absorb the excess liquid, creating a thin film, followed by rapid freezing of the grid. This protocol has been shown to introduce views in some proteins; however, even in these cases, the timescale of 100 ms has been large enough to allow the formation of denatured protein films on the surface (Dandey et al., 2020; Jain et al., 2012; Wei et al., 2018). Other options include using support films, such as carbon and functionalized graphene, or tilting the stage to obtain alternate views (Aiyer et al., 2023; Lu et al., 2022; Noble, Wei, et al., 2018; Tan et al., 2018; Xu et al., 2021). Tilting the stage or use of carbon support film suffers from a decrease in signal/noise ratio, especially if the protein is small, as the contrast is decreased due to either the support film or thick ice while imaging at higher tilt angles and tilting is less commonly used. While carbon support films are more commonly used in samples such as ribosomes, viruses, and some ion channels (Baker et al., 2021; Hesketh et al., 2018; Tobiasson et al., 2022), graphene/graphene oxide is suitable for low-molecular-weight specimens. These techniques have been successful in alleviating orientation bias in a few cases but are not generally applicable. Additionally, there is now some image processing software that deals with the orientation bias in the data better than others because of the better weighing of the different projections during reconstruction, but the problem still persists (Ramírez-Aportela et al., 2022; Sorzano et al., 2021).

The most widely used approach in decreasing the preferred orientation of macromolecules is to add surfactants to the sample buffer prior to grid preparation (Chen et al., 2019, 2022; B. Li et al., 2021). This method is popular because it is simple and does not require any additional steps, techniques, or instruments. A large number of surfactants are available and can be used, but care must be taken to avoid disrupting the protein’s native structure in the process. Often, extensive trial and error is involved when surfactants are screened to overcome the preferential orientation problem. EMDB has at least 83 different soluble protein structures (retrieved in 2022), in which a detergent is added to the buffer. The most popular choice of detergent is non-ionic or zwitterionic. Recently, cationic or anionic detergents have been found to work for a few samples but their use is not generalized yet (B. Li et al., 2021). We also encountered the preferred orientation problem in many of our projects. We asked if an informed decision on the choice of grid freezing conditions can be made based on the properties of the protein, thus minimising the time spent screening many surfactants and specimen preparation conditions.

To achieve this goal, we tested some commonly used surfactants with different properties on a set of five proteins, CRP-pentamer, CRP-decamer, Catalase, PaaZ and Spike. In addition, we explored the effect of the presence of the histidine tag on Spike and β-galactosidase and physical factors such as the temperature during the sample application step on catalase and PaaZ. We also serendipitously observed the effect of grid hole dimensions of the holey carbon grid, on the orientation distribution of catalase and have discussed this briefly. Through this analysis, we identified factors that affect and determine the macromolecule behaviour on grids before freezing and studied their effects with a focus on the preferred orientation problem. This account highlights the factors that contribute to orientation bias and provides valuable information that can assist in achieving the optimal freezing conditions for any given macromolecule.

## 2. Results

### 2.1. Analysis of preferred views of selected macromolecules

A set of well-characterized proteins with varying molecular weights, shapes, symmetries, surface charge distributions, and known experimental structures were selected, to rule out any artefacts that may arise owing to the changing conditions of sample preparation. The test proteins used for the analysis include C-reactive protein (CRP) pentamer, CRP decamer, human erythrocyte catalase, SARS-CoV-2 spike protein, *E. coli.* PaaZ and *E. coli* β-galactosidase. Micrographs and 2D class averages of these datasets collected at the start of the study are shown in Figure 2.

**Figure 2:**
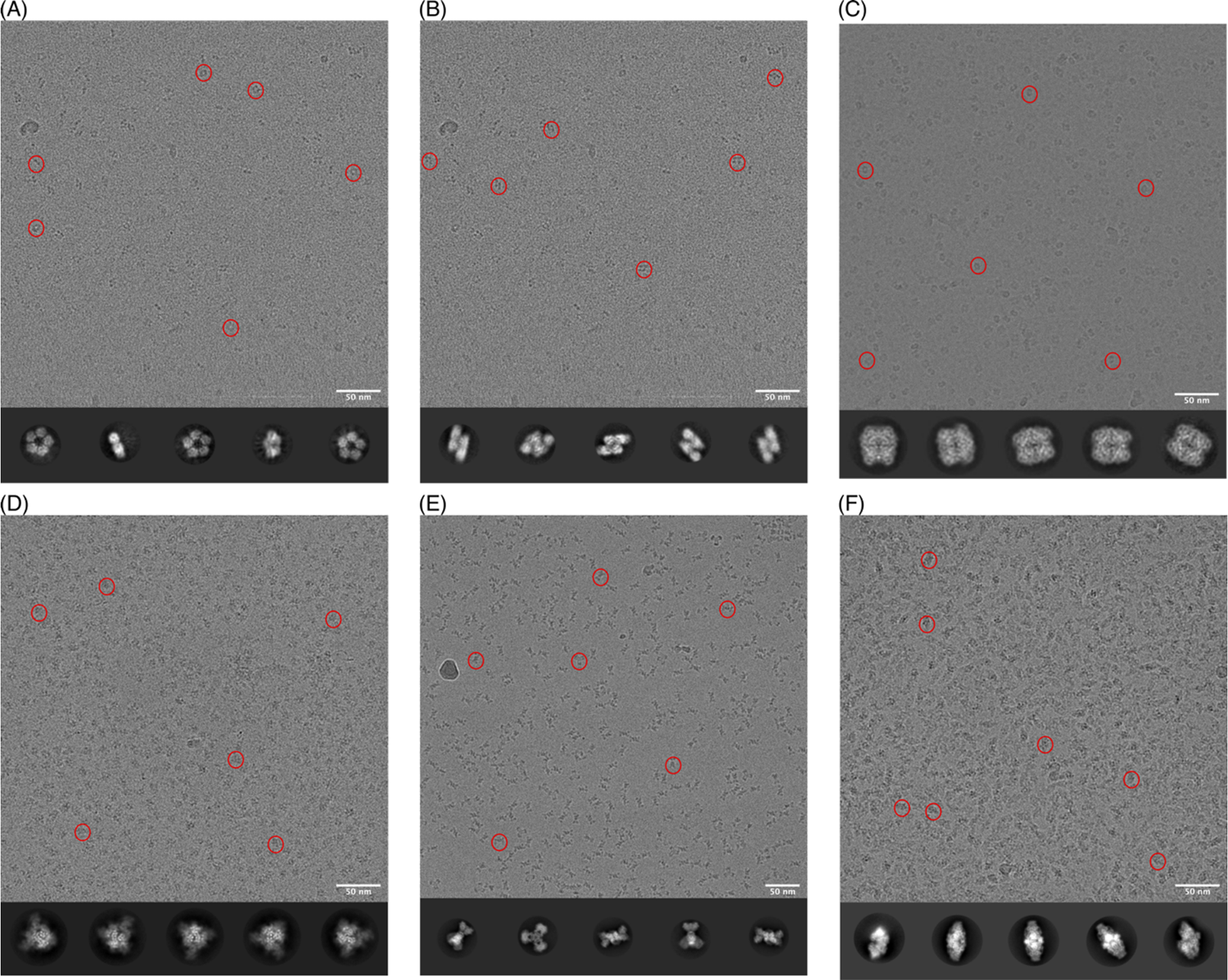
Representative micrographs with few select particles indicated with red circles and 2D class averages of test proteins used in this study. (A) C-reactive protein (CRP) pentamer adopts a preferred bottom view, which shows the pentameric arrangement of the monomers (B) C-reactive protein decamer adopts a preferred side view, which shows the staggered arrangement of two CRP pentamers stacked on top of each other. Same micrograph is used in A and B. (C) catalase adopts a preferred top view as seen in the micrograph and 2D class averages (D) SARS-CoV2-spike adopts a preferred bottom view showing the trimeric arrangement (E) PaaZ adopts a preferred side view as seen in the 2D class averages and the micrograph shows occasional clumping of hexamers on the grids (F) β-galactosidase with an N-terminal his tag adopts a preferred side view as seen in the 2D class averages and the micrograph shows aggregation on grids. Among the above datasets catalase and PaaZ grids were prepared at 4°C and all other grids were prepared at 20°C.

CRP exists in a concentration-dependent equilibrium of pentamer and decamer of a 25 kDa polypeptide under physiological conditions (Okemefuna et al., 2010). At the concentration range used for cryo-EM experiments, both pentamer and decamer populations were observed in the same micrograph and analyzed separately (Figure 2A, B). The effect that both these populations can have on each other’s behaviour in the thin film is an interesting concept that has not been analysed in the current study but is discussed briefly. C5 symmetry was applied to the smaller pentameric molecule, and no symmetry was applied to the decamer during image processing. The preferred view in the CRP pentamer is the top/bottom view, where the symmetric arrangement of the monomers can be seen. Some side views are observed, but no tilted side views are seen in the 2D class averages (Figure 2A). In contrast, in the case of the CRP decamer side views are predominantly observed and no top/bottom views are seen (Figure 2B). Catalase exists as a dimer of tetramers and possesses D2 symmetry (Ko et al., 2000). It adopts a preferred top/bottom view on the grids when the grids are prepared at 4°C using Quantifoil 0.6/1 holey carbon grids (Figure 2C). This enzyme has previously been noted for its peculiar behaviour during cryo-EM grid preparation and has been used as a test specimen to overcome orientation bias, and we used it as a control sample and to probe additional parameters (S. Chen et al., 2022; Fan et al., 2021; Vinothkumar & Henderson, 2016). The SARS-CoV-2 spike protein is now a well-studied trimeric viral membrane protein whose symmetry can vary depending on the conformation of the receptor-binding domain (Huang et al., 2020; Wrapp et al., 2020). Spike trimer has three receptor binding domains (RBD), one per monomer, and they can adopt different conformational states and thus dictate the symmetry of the protein. Soluble spike protein shows a preference for a bottom view on grids in the absence of an additive (Figure 2D). In this data set, we observed that there was heterogeneity in the RBD domain, and hence C1 symmetry was applied for the rest of the analysis.

PaaZ is a bifunctional enzyme and consists of six polypeptides, where a domain-swapped dimer assembles to form a trimeric structure. In an earlier study, when frozen in ice, the enzyme was found to form clumps, and aggregates, which was overcome by the use of graphene oxide and a low concentration of protein (Sathyanarayanan et al., 2019). PaaZ adopts a preferred side view in the absence of an additive when frozen in ice at 4°C, with a tendency to clump or cluster (Figure 2E). Nevertheless, the isolated particles were sufficient to obtain reasonable maps, and here we were interested in screening additional parameters to check if some of the aggregation and clumping could be overcome to enhance the quality of data of the enzyme in ice. β-galactosidase from *E. coli* is one of the well-studied enzymes as well as a test specimen in cryo-EM (Bartesaghi et al., 2018; Juers et al., 2012). It is a glycoside hydrolase enzyme, similar to catalase and exists as a tetramer with D2 symmetry. When purified with a tag at its N-terminus, the enzyme adopts a preferred side view and tends to form clumps or aggregates in ice (Figure 2F). Among all the standard proteins tested, spike, PaaZ and β-galactosidase were obtained by overexpression and had additional poly-histidine affinity tags at either the N or C termini of their monomers.

### 2.2. Surfactants affect macromolecule orientation distributions in a charge-dependent manner

We tested a specific set of surface-active molecules with varying head group charges (cationic, anionic, and non-ionic), chain length packing (unsaturated vs. saturated alkyl chain), differing in their critical micelle concentration (CMC) and concentration on the set of proteins described above. All the surfactants were used at concentrations lower than their respective CMCs. The properties of the surfactants are listed in Table 1.

**Table 1.**
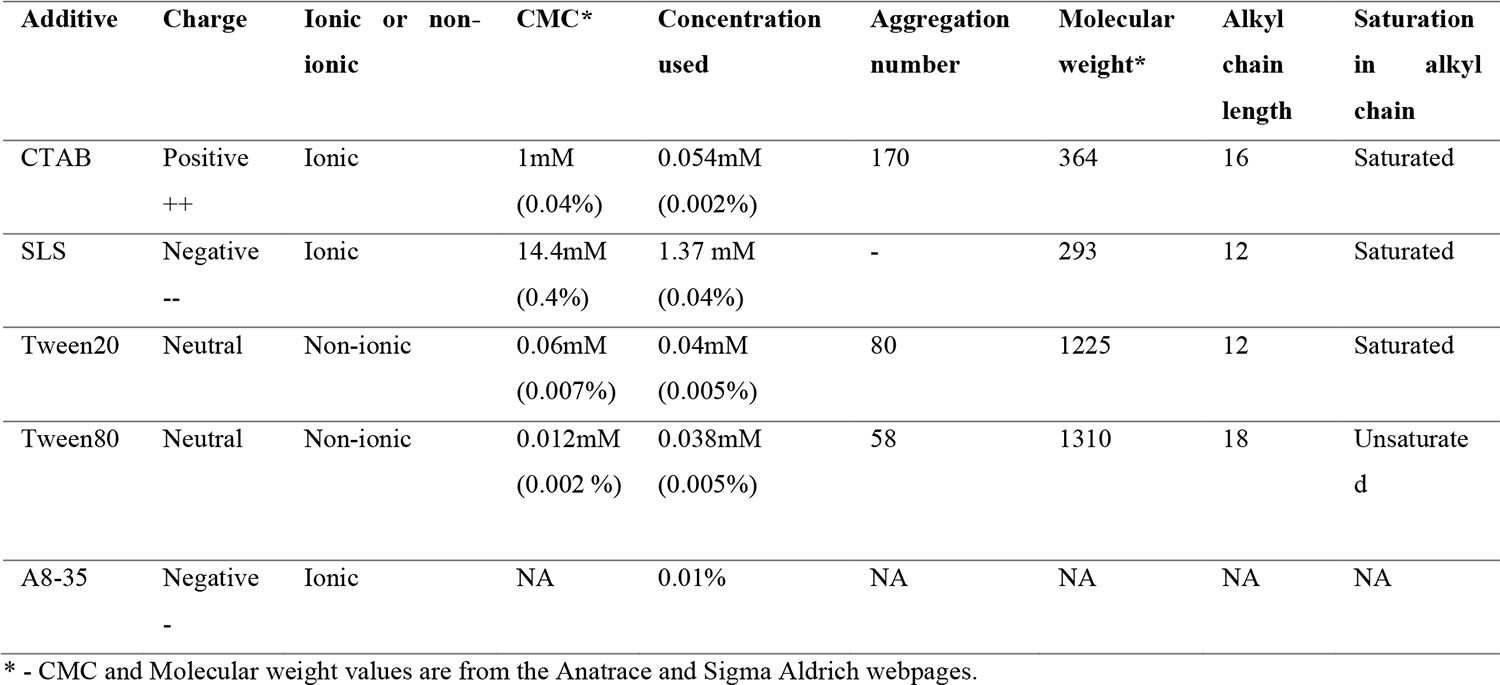
Properties of surfactants used in this study.

Upon the addition of cationic surfactant cetyltrimethylammonium bromide (CTAB), a change in the orientation distribution was observed for all the samples tested (Figure 3). An improvement in the orientation distribution of CRP pentamer and decamer was observed resulting in isotropic 3D reconstructions (Figure 3 A, B & Table 2). For catalase, the initial no-additive dataset was collected from a specimen prepared at 4°C on a Quantifoil 0.6/1 grid, which has a severe preference for the top/bottom views (Figure 2C). However, as the grids for all the additive datasets were made at 20°C, for an appropriate comparison, we prepared catalase grids at 20°C with no additive also. This dataset shows a preference for a tilted side view, annotated as side view A (Figure S2). The changes in orientation distributions of catalase are a result of variations in the incubation temperature and grid hole geometry, which we did not anticipate to have such a large effect on the quality of the reconstruction and were explored further in this study (discussed below). Upon addition of CTAB to catalase grids, the preference for the tilted side view A was lost, instead the particles adopted a preference for the other tilted side view B, and a map of similar quality to the no additive dataset was obtained (Figure 3C & Table2). In the case of PaaZ, the preference for the side view was maintained even in the presence of CTAB, but additional side-tilted views were sampled, which led to a better resolution and higher-quality map (Figure 3D & Table 2). The addition of a negatively charged detergent sodium lauryl sarcosine (SLS), resulted in evenly distributed orientation distributions, with no preference for any particular view for all cases tested except PaaZ (Figure 3). The resulting maps with SLS as additive in all the cases are of lower resolution, which may be due to a lower number of particles or the ice thickness and possible effect of the anionic headgroup (Table 2).

**Table 2.**
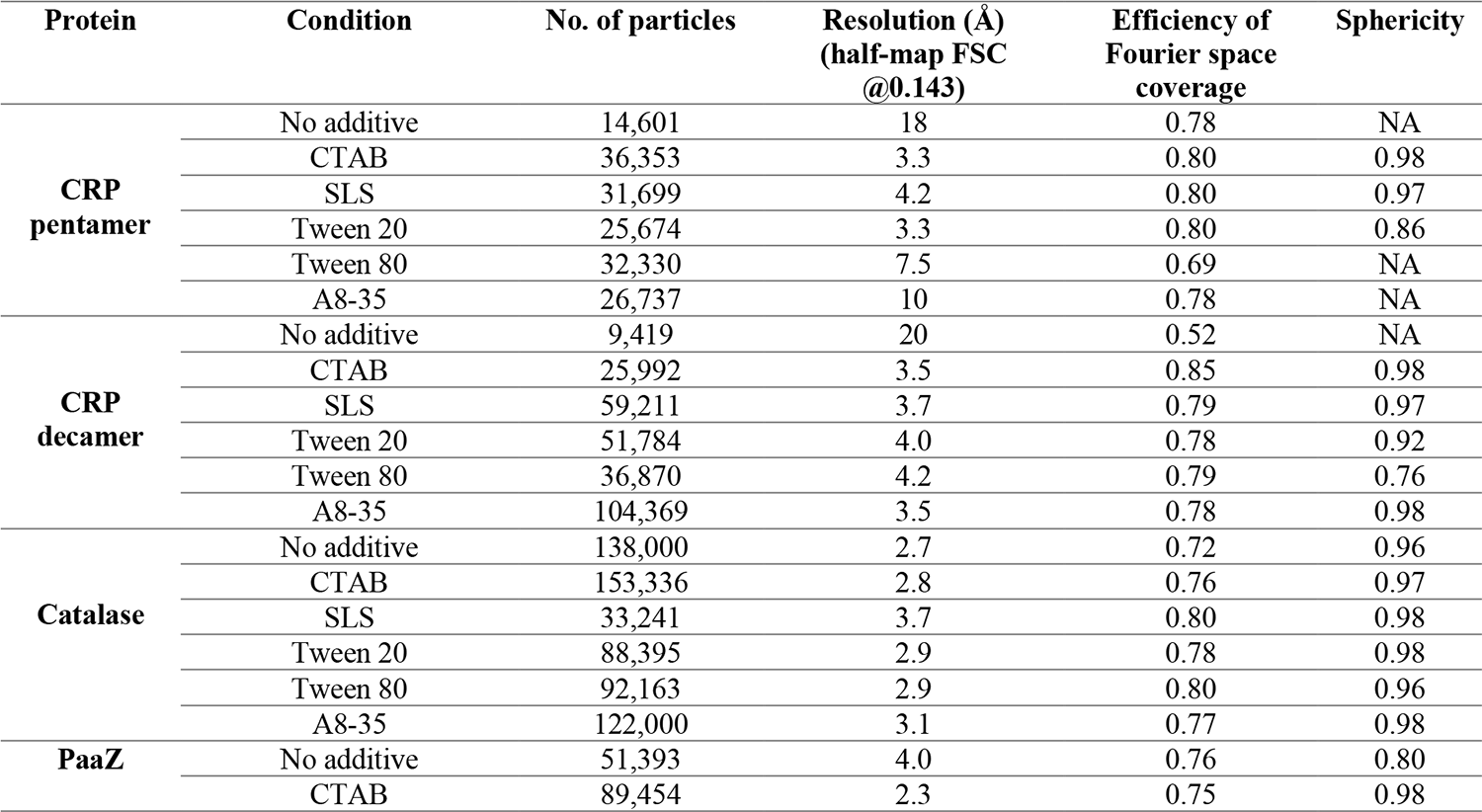
Comparison of parameters of the no additive and surfactant additive datasets.

**Figure 3:**
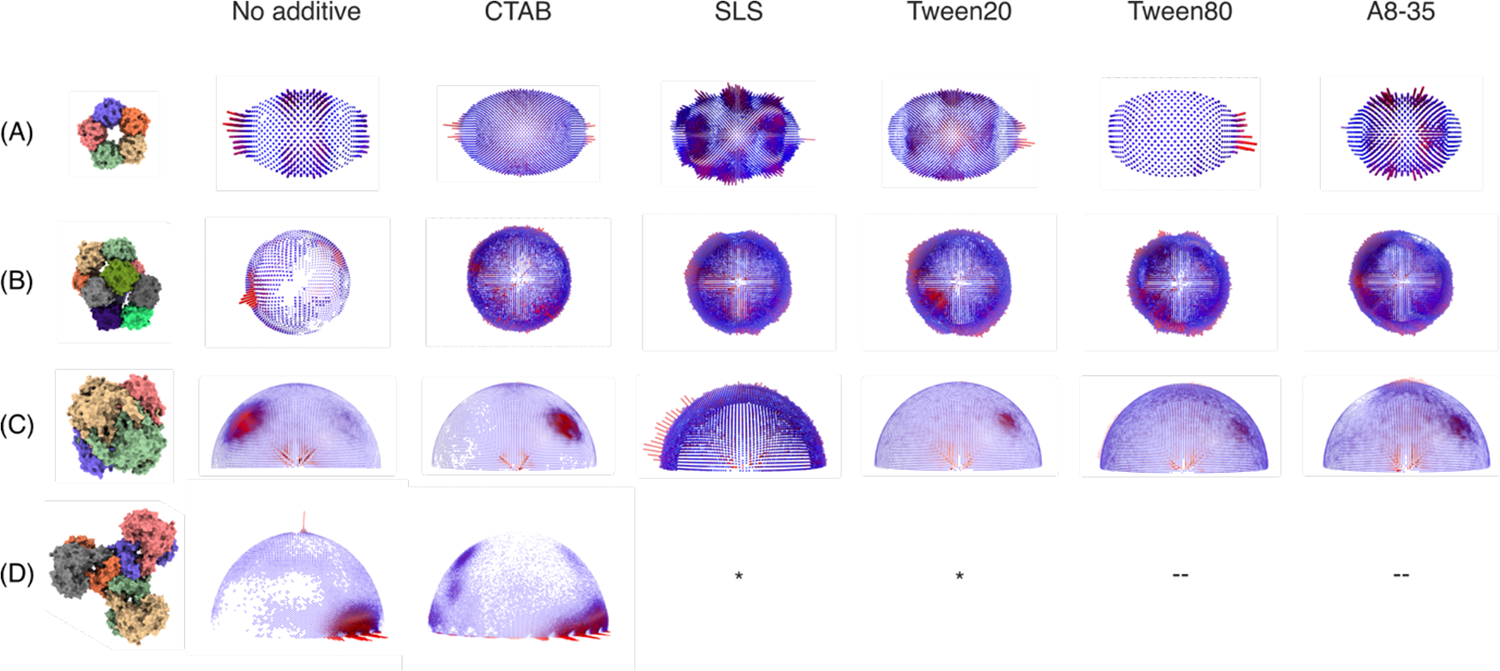
Orientation distribution plots from Relion (Scheres, 2012) of proteins upon the addition of surfactants with varying properties to the sample buffer before grid preparation. The reference structures of the respective proteins are generated by creating a surface representation in ChimeraX from models from PDB with IDs 7pkb, 7pk9, 1dgf, and 6jql. (A) CRP-pentamer orientation distribution change upon the addition of any surfactant. The distributions are distinct from each other except Tween 20 and Tween 80, which have similar distributions (B) CRP-decamer orientation distribution change upon surfactant addition, all surfactants lead to a similar even orientation distribution (C) catalase orientation distribution changes upon surfactant addition where the charged surfactants have distinct distributions (CTAB, SLS) and the neutral surfactants (Tween 20/80) and A8-35 show similar distributions (D) PaaZ orientation distributions change upon the addition of cationic surfactant CTAB. * The effect of SLS and Tween 20 on PaaZ was also tested, but visual inspection of the micrographs (Figure S3) showed no improvement and no data was collected and thus not included. -- The effect of Tween 80 and A8-35 was not tested on PaaZ.

The addition of non-ionic detergents Tween 20 and Tween 80 to CRP pentamer, decamer, and catalase led to evenly distributed orientations (Figure 3), but PaaZ showed aggregation on grids (Figure S3). Further, Tween 80 was tested as it is known to form denser layers at AWI compared to Tween 20 due to differences in the alkyl chain (length and unsaturation) (Szymczyk et al., 2018). Additionally, it has been shown to differ in its ability to separate protein films from the AWI as compared to Tween 20 (Rabe et al., 2020). In the macromolecules tested in our study, no significant difference was observed between the orientation distributions obtained from either of the Tween surfactant datasets (Figure 3). However, we note that with CRP pentamer, the orientation distributions look similar with both Tween 20 and 80 but only the addition of Tween 20 led to an isotropic map (R=3.3 Å), while the addition of Tween 80 led to a low-resolution map (R=7.5 Å). These differences can be a result of variations in ice thickness or due to differential interaction of Tween 80 with CRP pentamer and more experiments need to be performed to determine the cause of this discrepancy. Amphipol A8-35, a surface-active ionic polymer that does not form typical micelles and is thus different from other surfactants was also tested to see its effect on orientations. The orientation distributions obtained in this case were similar to those of the Tween 20 and Tween 80 datasets (Figure 3). The observed results with the given test samples indicate that the charge on the detergent headgroup is an important parameter, whereas the chain length and saturation of the hydrophobic chain have very little effect in modulating the orientation distribution of the macromolecules.

To understand the importance of the discrete views that were sampled and their contribution to the quality of the map, the efficiency of the orientation distribution (E_od_) using CryoEF (Naydenova& Russo, 2017) and the sphericity of the final maps using the 3DFSC (Tan et al., 2018) server were calculated (Table 2). E_od_ provides an estimate of the coverage of the 3D Fourier space from the 2D projections in the data, whereas sphericity provides a measure of anisotropy in the final maps by estimating the directional resolution. Among all the datasets, the CTAB datasets stand out and result in higher-resolution maps. We think this could be a result of optimum ice thickness, along with a sampling of the orientations that contribute more to the reconstruction, which is also demonstrated in Figure S1 by using a subset of particles from catalase data as an example.

### 2.3. The presence of solvent-exposed poly-histidine tag affects protein orientations in thin films

SARS-CoV-2 spike protein and *E.coli.* β-galactosidase are two well-studied samples by cryo-EM and multiple structures have been reported in the past (Bartesaghi et al., 2018; Bodakuntla et al., 2023; Esfahani et al., 2024; Hardenbrook & Zhang, 2022; Harvey et al., 2021; Juers et al., 2012; Wrobel, 2023). However, we observed orientation bias with these two recombinantly purified proteins, with differing severity. The spike protein showed severe orientation bias, which resulted in anisotropic maps of poor quality that could not be improved by surfactant addition (data not shown). β-galactosidase showed comparatively less orientation bias and led to a reconstruction of moderate quality but the map was still anisotropic (Figure 4B). Given that both of these proteins feature poly-histidine affinity tags, and considering that the preferred view in both cases may involve an exposed tag, we hypothesized that this tag could be inducing a bias in orientation (Figure 4A).

**Figure 4:**
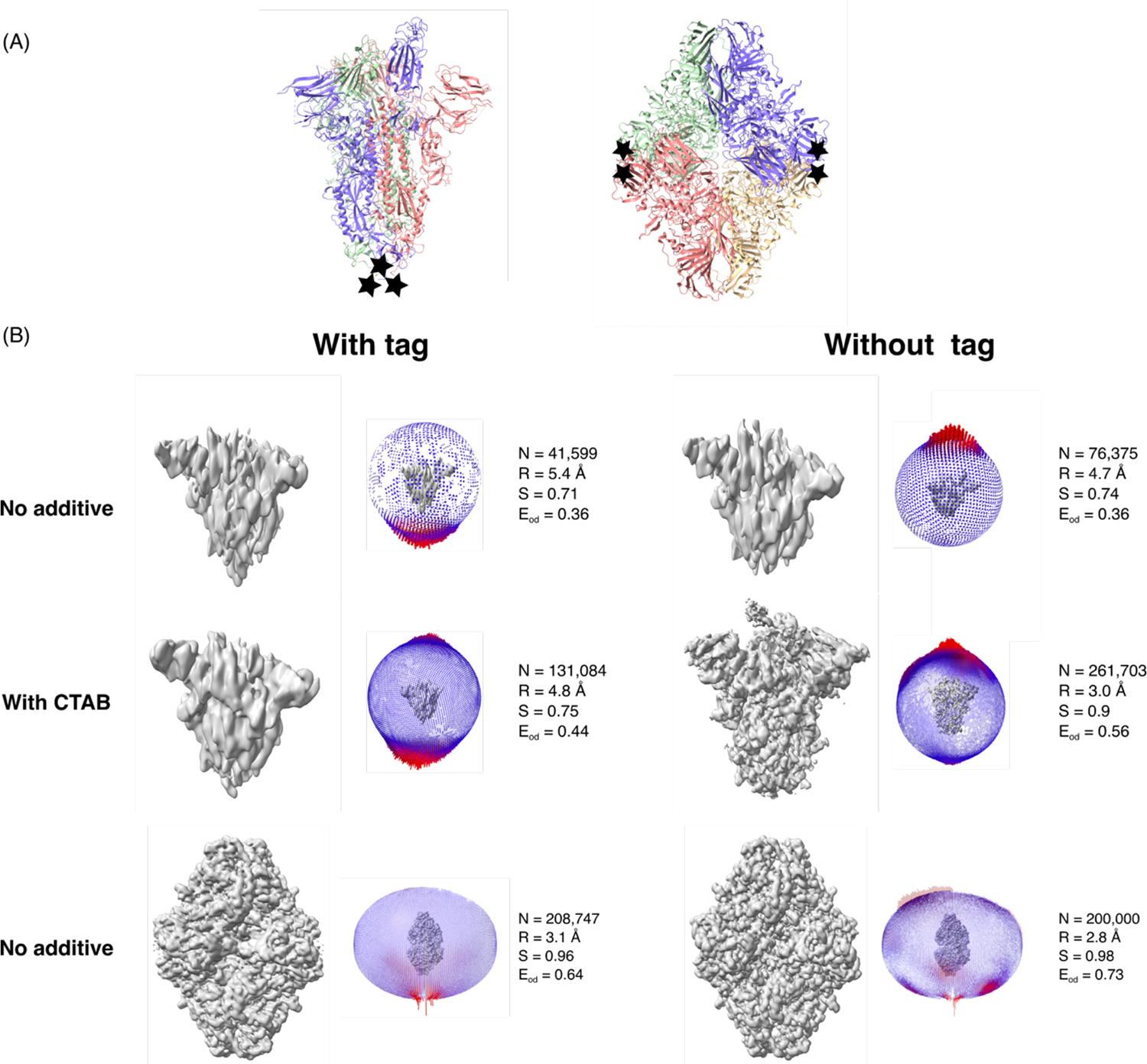
Effect of poly-histidine affinity tag on SARS-Cov2 spike protein and β-galactosidase orientation distribution. Different parameters that are used to analyse the quality of the maps are shown next to the orientation plots. N indicates the number of particles used for reconstruction, R indicates the final resolution of the map, S indicates the sphericity and Eod indicates the efficiency of Fourier space coverage. (A) The location of the tags on the protein models is indicated by black stars. The models used as reference are PDBs 8h3d and 6cvm for the spike protein and β-galactosidase respectively. (B) Orientation distribution plots of spike protein change upon removal of the affinity tag but the change is not enough to obtain an isotropic map. The addition of cationic surfactant CTAB further alters orientations of the spike protein without tag and leads to a more isotropic map. β-galactosidase enzyme orientations change upon removal of the affinity tag and lead to an isotropic high-resolution map without any additive. The unsharpened final combined maps are shown in grey in panel B.

Thus, the affinity tag from the spike protein and β-galactosidase were removed by protease cleavage, which led to a change in orientation distributions in both cases (Figure 4). In the case of spike, although the orientation bias remained, the preference shifted from the bottom view (where the histidine tag is attached) to the top view (Figure 4B). Subsequently, CTAB was added to the sample, imaging was performed on thin ice, and side-tilted views were obtained, which resulted in a reasonable isotropic map. The final reconstruction was obtained with 261,703 particles to an overall resolution of 3 Å (Figure 4B). At this resolution, amino acid side chains and glycosylation on the protein surface were visible in the ordered regions but the regions of RBD were poorly resolved substantiated by the local resolution plot (Figure S4). Further, we used the spike data set to evaluate the effect of different post-processing methods and the fit of the model to the map (He et al., 2023; Pintilie et al., 2020; Sanchez-Garcia et al., 2021)(summarized in Figure S4). In comparison to the other datasets, the increase in E_od_ and sphericity is evident in this case (Figure 4B). To verify that the improvement in map quality is a result of new orientations and not an increase in the total number of particles, all the top/bottom views were removed for 3D refinement, and the reconstruction using only the side/tilted views corresponding to ∼94,000 particles was performed. This resulted in a map with a sphericity of 0.88 and a resolution of 3.4 Å (Figure S5) similar to that with all the particles, illustrating that the top/bottom views contribute little information to the final reconstruction of spike protein.

In the case of β-galactosidase, the orientation bias was reduced and alternate views were sampled as indicated by the orientation plot and improvement of the E_od_ value from 0.64 to 0.73 when the tag is cleaved. The resulting reconstruction also shows improvement as indicated by visual inspection of the map (Figure 4B and Figure S6) and the improvement in resolution and sphericity. Zoomed-in areas of the map to highlight the difference in map quality and comparison of the half map and model vs. map FSCs of the β-galactosidase are shown in Figure S6.

### 2.4. The temperature of the incubation chamber during freezing affects protein orientations

The temperature at which the grids are held and blotted is critical for determining ice thickness, protein stability, and dynamics. Hence, grids were prepared at different temperatures to determine the effect of temperature on the orientation distributions.

Typically, lower temperatures are used during sample application to grids to keep the protein stable (except in some cases, such as microtubules, where a higher temperature is a prerequisite), and the temperature as a parameter is not very often screened to overcome the preferred orientations. For catalase, the orientation distributions varied significantly when grids were prepared at different temperatures. The coverage of the 3D Fourier space (E_od_) improved from 0.64 at 4°C to 0.72 at 20°C to 0.77 at 37°C (Figure 5A). However, the resolution of the final reconstruction is best for the 4°C dataset, which could be due to the higher number of particles used for reconstruction, that is, 478,656 compared to 138,031 for the 20°C dataset and 75,312 for the 37°C dataset. The respective B factors (from Relion post-process) for the reconstructions were −107 Å^2^, −110 Å^2^, and −126 Å^2^.

**Figure 5:**
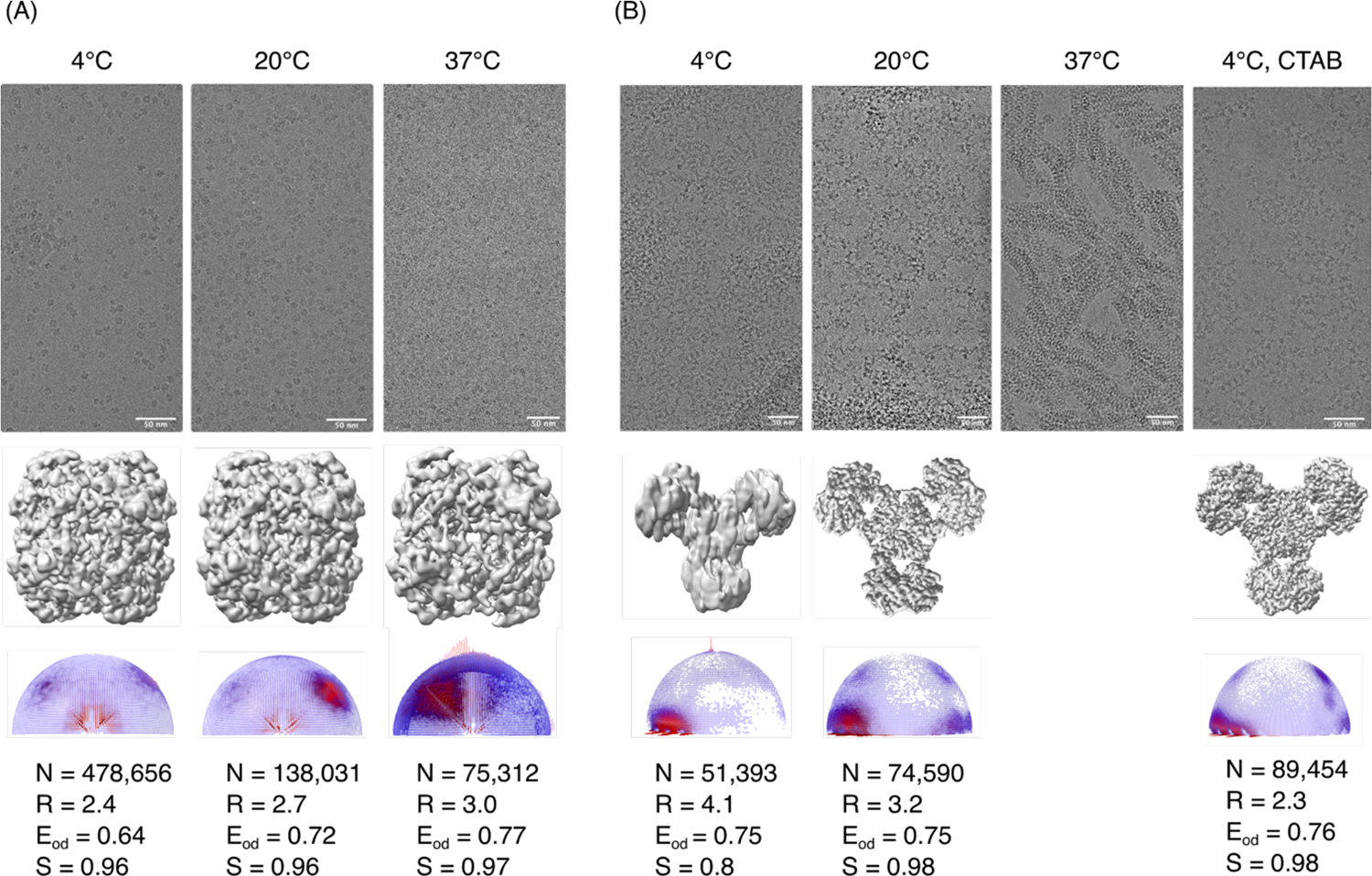
Effect of temperature during cryo-EM sample preparation of catalase and PaaZ. Micrographs, maps, orientation distribution plots and different parameters that are used to analyse the quality of the maps are shown. N indicates the number of particles used for reconstruction, R indicates the final resolution of the map, S indicates the sphericity and Eod indicates the efficiency of Fourier space coverage. (A) Catalase orientation distributions change significantly when grids are blotted at different temperatures in the absence of any additive (B) PaaZ orientation distributions change slightly when grids are held and blotted at different temperatures in the absence of any additive. In the case of PaaZ, the condition with grids prepared at 4°C with CTAB additive is included for comparison as this combination led to a high-resolution isotropic map.

In the case of PaaZ, when the grids were prepared at 4°C, 20°C and 37°C, increasing higher-order structures were observed, particularly at 37°C. At 4°C, higher-order structures were minimal, but a preference for the side view was observed. Data collected at 4°C and 20°C have similar E_od_, and the orientation distribution plots show a slight increase in the sampling of additional side-tilted views at 20°C, indicated by the reduction in empty spaces in the orientation plots obtained from Relion (Figure 5B). However, clumping of the particles was still observed at 20 °C. Hence, CTAB was added to the sample and the grids were prepared at 4°C, which led to less clumping and consequently resulted in a high-resolution map at 2.3 Å, described further in the next section.

It is evident from these observations that physical factors, such as the grid preparation temperature can affect protein behaviour and should be considered as an important screening condition when dealing with orientation bias along with surfactants.

### 2.5. High-resolution map of *E. coli* PaaZ in ice

To understand the mechanism of action of a macromolecule, high-resolution maps are required to build an accurate model and more importantly, to model the solvent molecules and other ligands of interest. We previously reported the structure of PaaZ, a bifunctional enzyme from *E. coli* that catalyzes two steps of the phenylacetic acid degradation (paa) pathway. In the previous study, the structure of PaaZ was determined at ∼2.9 Å using graphene-oxide support grids (Sathyanarayanan et al., 2019). During that study, it was observed that the enzyme tended to clump when frozen in ice in the absence of any additive, and the use of a graphene oxide support and a low concentration of enzyme yielded a very good distribution of particles. In the current study, we obtained a high-resolution map using CTAB as an additive during grid preparation, with D3 symmetry and higher-order aberration correction during image processing. The resolution estimated by comparison of the half maps FSC (at 0.143 threshold) is 2.3 Å, and a comparable resolution of 2.4 Å is obtained with the comparison of the map vs. model FSC (at 0.5 threshold) (Figure 6A). The model-fitted PaaZ structure is shown in Figure 6B. Several water molecules could be modelled at this resolution and an electrostatic potential map of the dimer with water molecules modelled as spheres and coloured in cyan is shown (Figure 6C). To further analyse the quality of the data, the Reslog plot was calculated by obtaining a 3D reconstruction using different numbers of particles and plotting the resolution (1/d^2^) obtained with the number of particles (lnN) used (Figure 6D) (Rosenthal & Henderson, 2003). From the experimental data, it can be seen that as the resolution approaches the Nyquist limit of 2.1 Å, the addition of more particles from 40,000 to 80,000 does not lead to an improvement in resolution. The slope of this graph was used to calculate a B-factor of 69.4 Å^2^, which is slightly lower than the B-factor estimated from the post-processing option in Relion, which was used for map sharpening (−75 Å^2^). A 200-particle dataset was used as a base, and the particles required to obtain a certain resolution were calculated theoretically. These theoretical estimates are in agreement with the experimental values.

**Figure 6:**
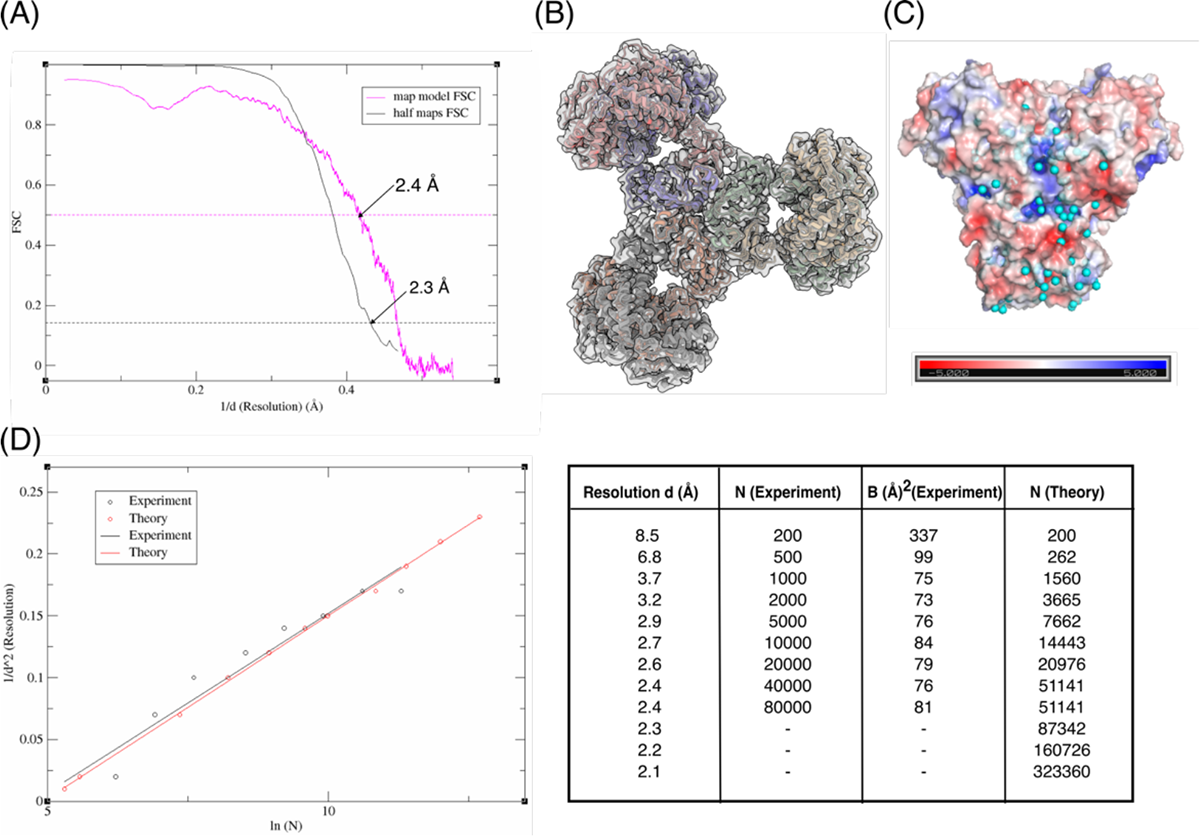
High-resolution Cryo-EM map of PaaZ grids prepared at 4°C with CTAB additive (A) Comparison of the half maps and map versus model FSCs of the PaaZ dataset. (B) The six polypeptides coloured individually and in cartoon representation fit into the cryo-EM map (transparent grey) of PaaZ. (C) Electrostatic potential surface representation of the domain-swapped PaaZ dimer with waters modelled and shown as cyan spheres. (D) Reslog plot of PaaZ with the experimental and theoretical numbers of particles required to reach a particular resolution. N indicates the number of particles used for reconstruction, d is the resolution, and B indicates the B factor, as estimated by the Relion post-processing. D3 symmetry was applied for the reconstruction, and the Reslog plot indicates the number of particles used, not the asymmetric units averaged.

## 3. Discussion

In this era of artificial intelligence-based protein structure prediction (Baek et al., 2021; Jumper et al., 2021; Schauperl & Denny, 2022), there is still no substitute for the joy one feels when the molecule of interest is imaged and observed for the first time after hard work of expression and purification. However, the first data collection for a new sample often does not result in a high-resolution structure, as there can be one or more challenges that need to be overcome. Of these, the preferred orientation of macromolecules and the possibility of denaturation due to exposure to AWI is a serious issue and one of the major bottlenecks. It is now known that the tendency of macromolecules to adopt preferential orientation when frozen in ice using cryo-EM grids is a consequence of protein-AWI interaction (J. Chen et al., 2019; Noble, Wei, et al., 2018), but the nature of this interaction remains unclear. On the other hand, amphipathic molecules have been used in cryo-EM grid preparation since the 1980s (Frederik et al., 1989). Their use in solving the preferred orientation of proteins by modulating the air-water interface has recently been appreciated in cryo-EM and is now being used more routinely for high-resolution structure determination (J. Chen et al., 2019; S. Chen et al., 2022; Li et al., 2021). At a fundamental level, several open questions remain: what causes proteins to adapt to certain preferred views? How do surfactants affect this behaviour? What is the role of other physical properties such as ice thickness, surface tension, and temperature in the behaviour of macromolecules during freezing? To address some of these questions, we systematically tested grid freezing conditions on a set of well-characterised proteins that adapt preferred orientations. The selected proteins had molecular weights ranging from 125 kDa to 466 kDa and varying symmetries (Figure 2). The degree of orientation bias also varied significantly. The CRP decamer and SARS-CoV-2 spike had a more severe orientation bias, with more than 90% abundance for the preferred view, whereas in the case of the CRP pentamer, the abundance was 50%, and in catalase, it was near 60%. Additionally, Spike, PaaZ and β-galactosidase were recombinantly overexpressed and had poly-histidine as the affinity tag at the termini of each monomer, whereas CRP and catalase, which were purified from native sources and did not contain any affinity tags.

The air-water interface exhibits an affinity for OH^-^ ions from water, resulting in a negative charge (Chaplin, 2009; Drzymala et al., 1999). Extensive research has been conducted on the behaviour of proteins and surfactants at the air-water interface and in thin films, spanning the fields of cryo-EM and surface chemistry (Li et al., 2021; Rabe et al., 2020; Samanta & Ghosh, 2011; Szymczyk et al., 2018). Based on these investigations and an examination of the surface charge distributions of macromolecules in our study, we postulated that electrostatic interactions may play a pivotal role in dictating protein orientations. To visualise the surface charge distributions of the proteins, electrostatic potential maps were calculated by assigning charges using the pdb2pqr server (Jurrus et al., 2018) at a given pH and visualised using ChimeraX (Pettersen et al., 2021). It was observed that the charge on the preferred view was positive in the case of catalase and neutral in CRP-pentamer, CRP-decamer, and PaaZ and negative and neutral in β-galactosidase (Figure S7). The preferred bottom view of the spike is negatively charged, but this charge is likely masked by the presence of a 56 amino acid-long affinity tag from each monomer. Further, the charge on the alternate view of β-galactosidase with tag, which is not seen in the 2D classes is more negative compared to the preferred view (Figure S7C).

An interesting sample in our study is the C-reactive protein (CRP) that exists as both pentamer and decamer populations dependent on concentration and, the impact of these populations on each other’s orientation distribution is intriguing. The particle numbers of each population of CRP and their relative abundance on the micrographs (after 2D classification) are summarized in Table S1. It’s plausible that one population exhibits a greater affinity for the AWI, monopolizing its free space and shielding the other from interaction. However, this scenario doesn’t seem applicable to CRP, as both pentamer and decamer populations adopt preferred orientations in the absence of additives indicating their interaction with the AWI. Furthermore, interactions and potential denaturation of a specific population at the AWI can alter the solution’s effective concentration, potentially causing a shift in the pentamer-decamer equilibrium. These factors are important in determining the behaviour of the protein on the grids but for this study, we have not considered these effects and analysed both the pentamer and decamer populations independently. The negatively charged B-face of CRP in the absence of additive is (shown in Figure S7B) less sampled in the case of the CRP pentamer and not sampled at all in the CRP decamer. When the surface charge at the AWI was changed to positive by the addition of cationic surfactant CTAB, the orientations changed in almost all cases, with a slight preference for certain alternate views (Figure 3). Hence, electrostatic interactions at the AWI are an important factor in determining protein orientations in the absence of any additive. When non-ionic surfactants Tween 20 and Tween 80 are added to the sample buffer, the observed orientation distributions are uniform (Figure 3). Among the non-ionic surfactants, the orientation plots are similar, as they all make the AWI charge neutral by occupying the free surface and work by diminishing the charge-based interactions between the protein and the AWI.

Although CTAB addition resulted in isotropic maps, the coverage of the 3D Fourier space was the least when compared with other surfactants (Table 2). The CTAB datasets showed a preference for one or more views, but in the macromolecules tested, these preferences were desirable. Upon addition of SLS, an even orientation distribution was observed for the CRP pentamer, CRP decamer, and catalase. Tween 20 and Tween 80 had similar effects on orientation, except in the case of the CRP pentamer. A8-35 had an intermediate orientation distribution, which led to isotropic maps (Figure 3). In the order of homogeneity of the orientation distributions, the surfactants can be ranked as SLS > Tween 20 = Tween 80 > A8-35 = CTAB. The map vs. model FSCs were calculated for the datasets to obtain an independent resolution estimate, and they closely match the resolution estimated by the comparison of the half-maps in most datasets (Figure S8 and Table S2). Further, the use of different surfactants or sample preparation at different temperatures does not affect the structure of catalase as shown by the density of heme in catalase (Figure S9).

Poly-histidine tag-dependent changes in orientation have been reported recently by Bromberg et al., 2022 but the mechanism by which this occurs remains unclear. The addition of poly-histidine tag is a convenient method for expressing and isolating proteins (Hochuli et al., 1988; Merchant et al., 1998). Unlike crystallography, it is intuitive to think that the flexibility of histidine tags does not pose a problem in structure determination by cryo-EM as the molecules are imaged individually, and the tag is averaged out during the reconstruction process if it is present in random conformations. At pH 8, which is more often used as the sample buffer, histidine should be deprotonated and exist as a hydrophilic polar group, and why this should cause severe preference in orientations is not clear to us. The microenvironment at the AWI is complex and could be interacting with the histidine tag favourably and stabilizing it in that orientation.

Among the samples studied here, spike, PaaZ and β-galactosidase had a poly-histidine tag at the termini of the monomers. While the presence of the tag had no detrimental effect on the orientation distribution and reconstruction of PaaZ, it affected the orientations of the spike and β-galactosidase proteins significantly, and the addition of surfactants was not sufficient to mitigate the bias in the case of spike. In the case of PaaZ, the presence of a poly-histidine tag also causes a preference for the view that contains the affinity tag but the side view of the protein which contributes maximally to the reconstruction and hence even with very little sampling of alternate views the reconstruction is not as anisotropic as spike. We note that the his-tag might cause the clumping observed in PaaZ and this remains to be tested. Further, the use of D3 symmetry in PaaZ and D2 symmetry in β-galactosidase as compared to C1 symmetry in the spike contribute to the quality of the final reconstruction. As the spike protein and β-galactosidase have been extensively studied and many structures have been reported we wondered what could be the reason that we see such severe orientation bias and others do not. From the published structural studies on the SARS-CoV2 spike it was realised that there have been variations in the reported specimens used for cryo-EM in terms of the expression system, detergents added to the sample, cleavage of tags, buffers, etc (Bangaru et al., 2020; Bodakuntla et al., 2023; H. Cheng et al., 2023; Hardenbrook & Zhang, 2022; Wrapp et al., 2020; Wrobel, 2023). The variability in the spike protein’s behaviour is outlined in the 2022 review authored by Chua et al., 2022. They provide instances of the spike protein showing a preference for very thick ice (100 nm), displaying unfavourable behaviour on gold grids, and exhibiting notably low particle concentrations on grids in the absence of a detergent (Chua et al., 2022). Other parameters that might affect the orientation distribution include the glow discharge settings for grids that change the hydrophilicity of the grid surface, grid type (for example, Quantifoil with carbon support or Ultrafoil with gold support), grid hole diameter (0.6/1, 1.2/1.3, or 2/2), freezing temperature, and blotting duration. In our case, the removal of the affinity tag and CTAB addition to the sample buffer was required to sample other views of spike protein, which resulted in a map of reasonable resolution and improved isotropy (Figure 5B). In the case of β-galactosidase, the removal of the affinity tag was sufficient to obtain a high-resolution isotropic without the need for any additive (Figure 5B). We note that β-galactosidase with a C-terminal tag also showed preferred orientation in ice as recently reported by Esfahani et al., 2024. Thus, the effect of the poly-histidine tag is a key factor to be tested when faced with orientation bias.

Furthermore, physical factors, such as the temperature of grid freezing and grid type, can affect the mechanics of thin film formation and evaporation/blotting and, in turn, influence protein behaviour. We show that the temperature of grid freezing can change the orientation distributions of proteins by affecting the ice thickness and protein-protein interactions (Figure 5). Additionally, varying only the hole diameter in a holey grid and keeping other factors constant led to a change in the preference of views in the case of catalase (Figure S2). In Quantifoil 1.2/1.3 grids, the approximate ratio of the area occupied by the hole to carbon is 38:62%, whereas, in 0.6/1 grids, the ratio decreases to 30:70%. This change can affect protein distribution in the holes, carbon, and AWI. It is now evident that this also significantly affects the orientation distribution of macromolecules in the thin films formed in the holes. We note two caveats in this study: a) all the grids used were from Quantifoil with amorphous carbon as the support film, and it is unclear if similar behaviour will be observed with C-flat grids from Protochips that are manufactured differently or other grids such as UltraFoil or NiTi alloy (Fan et al., 2021; Quispe et al., 2007; Russo & Passmore, 2016; Schürmann et al., 2017). b) only one sample (catalase) was tested and further extensive study with more samples is required to generalise the effect of the holey grid geometry.

Previous studies have used surfactants to determine the structures of the proteins examined in this study. A comparison of these results with our observations is presented below. Catalase has been used as a standard test sample in the context of the preferred orientation problem in two independent studies, which take different approaches to solve the problem. In an earlier study, several high-CMC detergents were screened, among which CHAPSO resulted in the best orientation distribution and map quality, although a high protein concentration of 30-40 mg/ml was used to obtain a good distribution of particles and a small data set was sufficient to obtain a 2.2 Å reconstruction by averaging 119,000 particles with an E_od_ of 0.76 (S.Chen et al., 2022). Furthermore, the effects of ionic strength and pH were tested and were observed to have little or no effect on the protein orientation (S.Chen et al., 2022). Our results, in extension to this study, provide an understanding of the effects of surfactant headgroup charge, temperature of grid freezing, and hole carbon geometry on the orientations of catalase (Figure 3, 5 & S2). In another study, nickel/titanium grids coated with 2D crystals of the hydrophobic protein HFBI (which shields the protein from its exposure to AWI) and 2.3 mg/ml of catalase were used for preparing specimen and subsequent data collection and structure determination (Fan et al., 2021). The resulting catalase map has a resolution of 2.3 Å from 169,897 particles with an E_od_ of 0.80, which is comparable to our datasets (Table 2).

Similarly, multiple structures of CRP pentamer and decamer were reported in a study by (Noone et al., 2021), which focused on the effects of pH and ligand addition on the complement binding properties of CRP. One of the grid freezing conditions reported was CRP apoprotein at pH 7.5 with 0.05% Tween 20 added and the grid held at 4 °C and 65% humidity. These freezing conditions are similar to those described here, except for the concentration of Tween 20 used and the temperature and humidity during freezing. The orientations observed in this case look similar, and the resulting reconstructions from 256,289 and 204,354 particles for the pentamer and decamer (C5 and C1 symmetry) resulted in resolutions of 3.2 Å and 2.8 Å, respectively. In comparison, we have obtained a 3.3 Å map for the pentamer and a 4 Å map for the decamer by averaging 25,000 and 51,000 particles, respectively (Table 2). In the field of cryo-EM, the biggest variation is observed in sample preparation, sometimes using the same instrument. Therefore, it is reassuring to observe that other independent investigations (along with different grid freezing procedures) of catalase and CRP yield 3D reconstructions of comparable quality.

The air–water interface is a complex environment; the properties of the bulk and the surface vary drastically, and it is an active area of research in surface chemistry and aerosol fields (Martins-Costa & Ruiz-López, 2023; Nguyen et al., 2020; Zhong et al., 2019). Only the effects that occur at the surface can be observed, and trends can be analysed to improve the sample preparation methods. The explanation as to why they occur cannot be understood using standard cryo-EM experiments alone and requires complementary approaches such as infrared spectroscopy or dynamic surface tension measurement studies (Carter-Fenk et al., 2021; F. Tang et al., 2020). Alternative approaches to sample preparation have been proposed in recent years to overcome the preferred orientation issue (Esfahani et al., 2024; Drulyte et al., 2018; Glaeser & Han, 2017; Jain et al., 2012). One such approach was recently reported by Huber et al., where the sample was filled with nanosized capillaries made of silicon-rich nitride membranes embedded on a chip that physically controls the ice thickness. No blotting is involved in this process, and some test samples have demonstrated the potential of this method (Huber et al., 2022). However, the air-water interface is replaced by the solid-water interface, which may contribute to some effects of its own. Further use and standardisation of such methods are required to understand their general applicability.

## 4. Conclusion and outlook

One of the goals of single particle cryo-EM is to collect a smaller data set and average minimum number of particles to obtain a high-resolution reconstruction (Henderson, 1995). However, in reality, many factors affect this including the sample heterogeneity, detectors, beam-induced motion etc., and the recently observed effect of AWI and preferred orientation can be added to this list. We performed a comprehensive examination of how macromolecule orientations respond to alterations in physical factors, such as freezing temperature, and chemical factors, such as the addition of surfactants or the presence of affinity tags, during grid freezing for a standard set of proteins. This analysis provides insights into the behaviour of proteins on grids and can be utilised to address the preferred orientation problem systemically for any given macromolecule. When using surfactants, it is crucial to carefully assess protein stability using other techniques such as native gel electrophoresis, differential scanning fluorimetry, or size-exclusion chromatography before data collection to save time and resources. Furthermore, our findings highlight the necessity to innovate and create small-molecule additives that are inert to biological samples and can effectively occupy the AWI and alleviate its deleterious effects on proteins.

## 5. Materials and Methods

### Source of proteins

Human C-reactive protein (Cat. No. C4063) and Human erythrocyte catalase (Cat. No. C3556) were obtained from Sigma Aldrich chemicals. The protein samples were either concentrated using an Amicon 100kDa concentrator or diluted in respective buffers for grid freezing. All the detergent stocks were made in ultrapure water and dilutions were made and used on the day of the experiment. PaaZ was expressed and purified as described in (Sathyanarayanan et al., 2019).

SARS-CoV-2 S plasmid was a kind gift from the Krammer laboratory at Icahn School of Medicine at Mount Sinai. The spike gene was amplified from the plasmid and subcloned in the BacMam vector with a C-terminus HRV-3C cleavage tag followed by a 7x poly-histidine and twin strep tag. Bacmid DNA and virus were prepared as described in the Invitrogen bac-to-bac manual. After two generations of amplification in Sf9 cells, the V2 virus was used for transfection of HEK293F cells at 2 million/ml density. Sodium butyrate (4 mM) was added to enhance the production of protein 8 hours post-infection. The media supernatant containing the secreted spike protein was harvested on day 3 by centrifuging the cells at 150 g for 10 minutes. The media was incubated with pre-equilibrated Ni-NTA beads (Qiagen) at room temperature for 1-2 hours (1 ml beads/200 ml media). Ni-NTA beads were washed with PBS buffer containing 20 mM imidazole followed by elution with 280 mM imidazole in PBS buffer. The eluted protein was run on SDS-PAGE to assess purity and further concentrated and injected on a 24 ml column Superdex 200 size exclusion to exchange the buffer to 50 mM Tris pH 8, 200 mM NaCl,1 mM DTT. To cleave the tag, the eluted fractions from Ni-NTA chromatography were diluted with 50 mM Tris pH 8, 200 mM NaCl, and 1 mM DTT buffer and incubated with HRV3C protease overnight at 4°C, followed by reverse IMAC to obtain the spike protein without tag in the flow through. The flow-through was concentrated using an Amicon 100 kDa concentrator and flash-frozen using liquid nitrogen and stored at −80°C until further use.

β-galactosidase with N-terminal 6X histidine tag followed by a thrombin cleavage tag was a gift from Prof. Doug Juers, which was cloned in the pET15b vector and was transformed in *E. coli.* Strain JM109 (Juers et al., 2000). The cells from glycerol stock were patched on an LB agar with ampicillin and allowed to grow overnight at 37°C. A single colony was picked and allowed to grow in LB with ampicillin overnight. The next day 4 litres of LB with 100 μg/ml ampicillin was inoculated and OD_600_ was monitored every 30 minutes. The expression of protein was induced by adding 1 mM IPTG when the OD_600_ of the cells reached 0.6 and the cells were allowed to grow for 4-5 hours at 37°C. Subsequently, cells were harvested by centrifugation at 4000g for 20 minutes and the cell pellet was flash-frozen and stored at −80°C until further use. On the day of purification, the cells were resuspended in ∼50 ml of resuspension buffer containing 20 mM Tris pH 8, 500 mM NaCl, 5 mM Imidazole, 1 mM β-mercaptoethanol (β-ME), 1 mM MgCl_2_ and sonicated for 5 minutes (5 secs on 10 secs off) at 40% amplitude. The lysate was clarified by centrifugation at 18000 g for 45 minutes and the supernatant was collected. A 5 ml Ni-NTA column was equilibrated with 5 CV resuspension buffer followed by sample application of the supernatant at a flow rate of 1 ml/min. After the binding of the protein, the column was washed with 50 CV of resuspension buffer followed by a linear gradient elution with buffer B that consisted of resuspension buffer with 500 mM imidazole and 1.8 ml fractions were collected. The fractions from Ni-NTA elution were analyzed using SDS-PAGE and the fractions from the last half of the peak were pooled and dialyzed with 2×4L of dialysis buffer containing 25 mM Tris pH 8, 125 mM NaCl, 2.5 mM CaCl_2_, 2 mM β-ME. The protein was concentrated using a 100 kDa cutoff Amicon concentrator, aliquoted and stored at −80° C. An aliquot of this protein was thawed and injected into a Superdex 200 column and the buffer was exchanged to 100 mM Tris pH8, 200 mM NaCl, 2.5 mM MgCl_2_, 5 mM CaCl_2_, 2 mM β-ME. The cleavage of the tag was carried out by thrombin protease at room temperature for 4 hours followed by reverse IMAC to collect the protein without tag in flow-through. The protein without tag was concentrated and grids were prepared.

### Grid preparation

6.3 µl of the protein was thawed on ice and 0.7 μl of 10 x additive (surfactant) stock was added to get a final concentration of 1 x. This sample was incubated on ice for 2-5 mins and then centrifuged at 21,000g for 20 mins. Meanwhile, Vitrobot Mark IV (ThermoFisher Scientific) chamber was equilibrated at 20° C (unless stated otherwise) and 100% humidity. Quantifoil 1.2/1.3 or Quantifoil 0.6/1 grids were glow discharged in a reduced air environment using a PELCO EasiGlow chamber using a standard setting of 25 mA current for 1 minute. The grid was mounted on the Vitrobot Mark IV (ThermoFisher Scientific) and 3 µl of the sample was applied to the grid. A blotting time of 3-4 seconds, a wait time of 10 seconds and a blot force of 0 were used to obtain a thin film of the specimen. For the datasets where grids were prepared at different temperatures, the protein was incubated on a thermal block at the required temperature for 3-7 minutes before applying it to the grid. The Vitrobot chamber was maintained at the required temperature and 100% humidity. The blot time, blot force and wait time were kept constant.

### Grid screening and data collection

Grids were screened on Titan Krios microscope operating at 300 kV using standard low-dose settings and the automated data collection was set up either on the Falcon 3 or Gatan K2 detectors in counting mode with EPU software (ThermoFisher Scientific). Magnification of 59,000 x was used only for the catalase 37 °C dataset with a pixel size of 1.38 Å (and PaaZ enzyme incubated at 37°C with no additive but no data was collected). For all the other datasets collected on Falcon3, a magnification of 75,000 x corresponding to a pixel size of 1.07 Å and a dose of approximately 1 e^-^/Å^2^/frame were used and movies of 24-25 frames were collected. The datasets on the K2 detector (Gatan) were collected in EFTEM mode with a slit width of 20 eV and at a nominal magnification of 130,000 x corresponding to a pixel size of 1.08 Å. The dose was 5 e^-^/p/s and an exposure of 8 seconds was used. Each movie was fractionated into 32 frames with a total dose of ∼36.28 e^-^/Å^2^. The grid preparation condition and data parameters are summarized in Table 3.

**Table 3.**
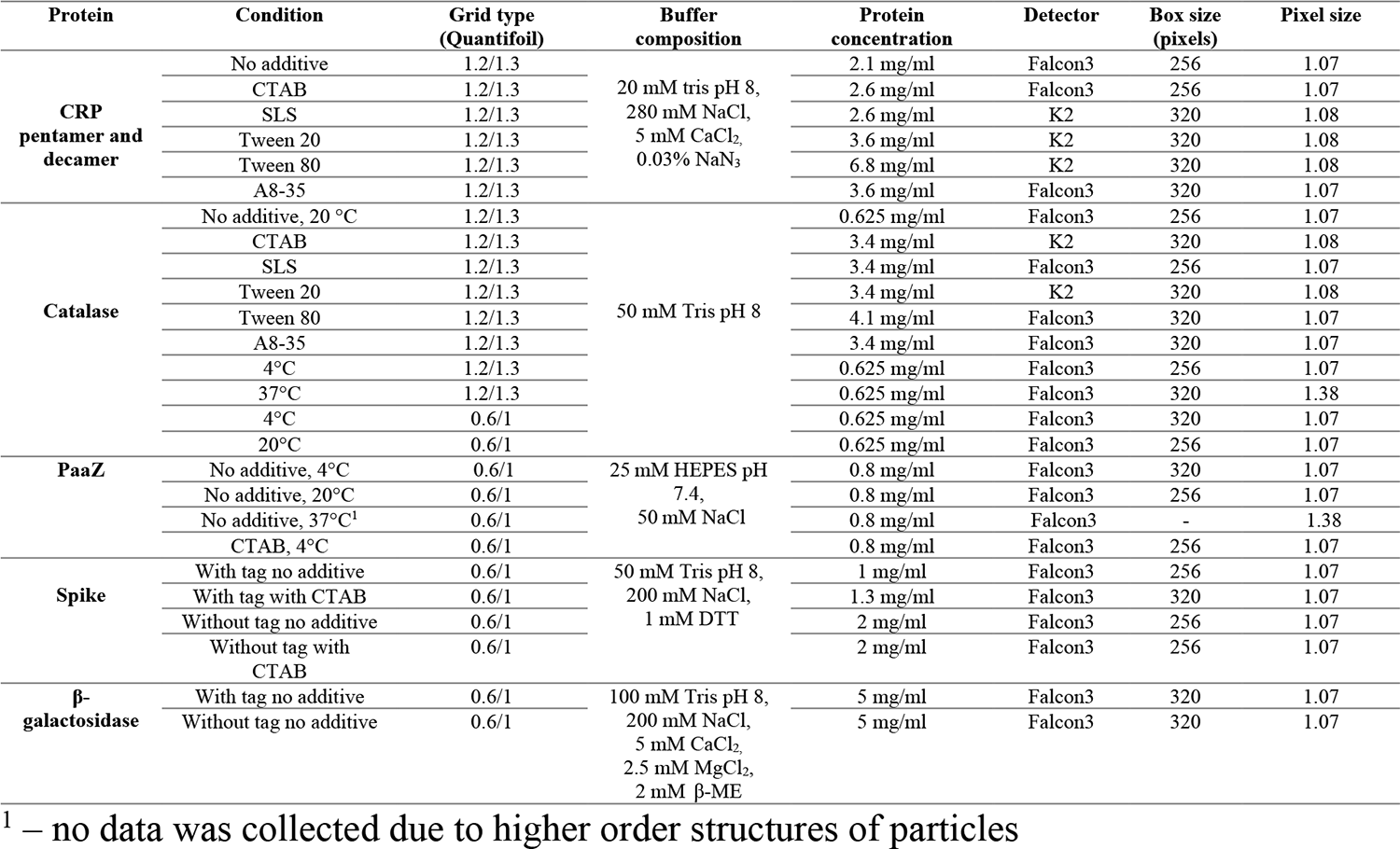
Summary of parameters of datasets collected under different conditions. All the imaging was performed in counting mode.

### Data processing and model refinement

RELION 3.1. (Scheres, 2012, Zivanov et al., 2018) was used to process all the data with a standard workflow which is summarized as follows. The multi-frame movies were summed and corrected for beam-induced motion using Relion’s in-built algorithm Motioncorr. The resulting summed micrograph images were used to estimate the contrast transfer function (CTF) using GCTF (Zhang, 2016). The micrographs after CTF correction were used to pick particles using either reference-free methods such as the LoG-based picking in Relion or Gautomatch (https://www2.mrc-lmb.cam.ac.uk/download/gautomatch-053/) or using previously obtained 2D templates. After picking, the particles were boxed and extracted using the box sizes of 256 or 320 pixels (Table 3). These particles were then subjected to multiple rounds of iterative 2D classifications to obtain good-quality, high-resolution classes. A low-resolution initial model was either generated from the same dataset or a model from pre-existing dataset was used as a reference for 3D refinement. The selected good-quality particles were then used for either 3D classification or 3D refinement based on the data (Scheres, 2012). If the resolution at this stage was below 4.5 Å, CTF refinement and Bayesian polishing were performed on the final particles to correct for beam tilt, anisotropic magnification, per-particle defocus and the effects of beam-induced motion respectively (Zivanov et al., 2018, 2019). Another round of 3D refinement was performed after this step. Further, postprocessing was done to sharpen the final map and estimate the final global resolution (Rosenthal & Henderson, 2003).

PaaZ-CTAB and Spike-CTAB datasets were subjected to multiple rounds of 3D classification. For most datasets (all except CRP pentamer), as a part of standard workflow the resulting map after reconstruction had the wrong hand and the maps were flipped to obtain the correct hand before the model was fit and refinement was performed. The 3D FSC server was used to calculate the sphericity of the final maps (Tan et al., 2018). The E_od_ was calculated using the cryoEF by using a subset of 1000 angles and the appropriate symmetries for respective molecules (Naydenova & Russo, 2017). ChimeraX (Pettersen et al., 2004) was used for generating the electrostatic potential maps of the structures. The Pymol APBS plugin was used to generate the PaaZ electrostatic potential map (Schrödinger, LLC, 2015). EMAN2 (G. Tang et al., 2007)and Phenix (Liebschner et al., 2019) were used to generate the map vs. model FSCs. The final model refinement was done in real space for the PaaZ and CRP datasets using Phenix (Afonine et al., 2018) and in reciprocal space using Refmac Servalcat (Murshudov et al., 2011; Yamashita et al., 2021) for catalase and spike datasets. Initial water picking in the PaaZ dataset was performed with Coot (Emsley & Cowtan, 2004) and manually inspected to remove those that were not hydrogen bonded to amino acid residues, also using the fofc map from servalcat (Yamashita et al., 2021) and a conservative approach was used so that noise was not modelled. Figures were made with Chimera and Pymol (Pettersen et al., 2004; Schrödinger, LLC, 2015).

## Acknowledgements

We acknowledge the National Cryo-EM facility, Bangalore, for data collection, which was supported by the Department of Biotechnology, DBT/PR12422/MED/31/287/2014, and the computing facility in the Bangalore Life Science Cluster. We thank Prof. Ramaswamy S and all the lab members for critical reading of the manuscript. SY acknowledges the graduate fellowship from TIFR/NCBS. KRV acknowledges the support of the Department of Atomic Energy, Government of India, Government of India, under Project Identification No. RTI4006. KRV is part of the EMBO Global Investigator Network. KRV acknowledges the discussion with Drs Pamela Williams and Judith Reeks, Astex, UK on β-galactosidase and the effect of tag.

Conflict of interest: The authors declare no conflict of interest.

## Data Availability

The cryo-EM maps and the coordinates have been deposited in the EMDB and PDB respectively with the following accession codes. CRP Pentamer with CTAB – EMD-37864 and PDB 8WV4 CRP decamer with CTAB – EMD-37865 and PDB ID 8WV5 PaaZ with CTAB at 4 °C – EMD-37866 and PDB ID 8WV6 Spike with CTAB – EMD-37953 and PDB ID 8WZI Catalase with CTAB – EMD-37956 and PDB ID 8WZM Catalase with SLS – EMD-37952 and PDB ID 8WZH Catalase at 4 °C – EMD-37955 and PDB ID 8WZK Catalase at 20 °C – EMD-37954 and PDB ID 8WZJ β-galactosidase, with tag – EMD-39809 β-galactosidase, with no tag – EMD-39808

## Notes

### Competing Interest Statement

The authors have declared no competing interest.

### Summary of Updates

Table 2 of surfactants has been updated. Figure 5 has been replaced with a revised version.

