## Supplementary_information for "Factors affecting macromolecule orientations in thin films formed in cryo-EM"

### S1.1. Supplementary information

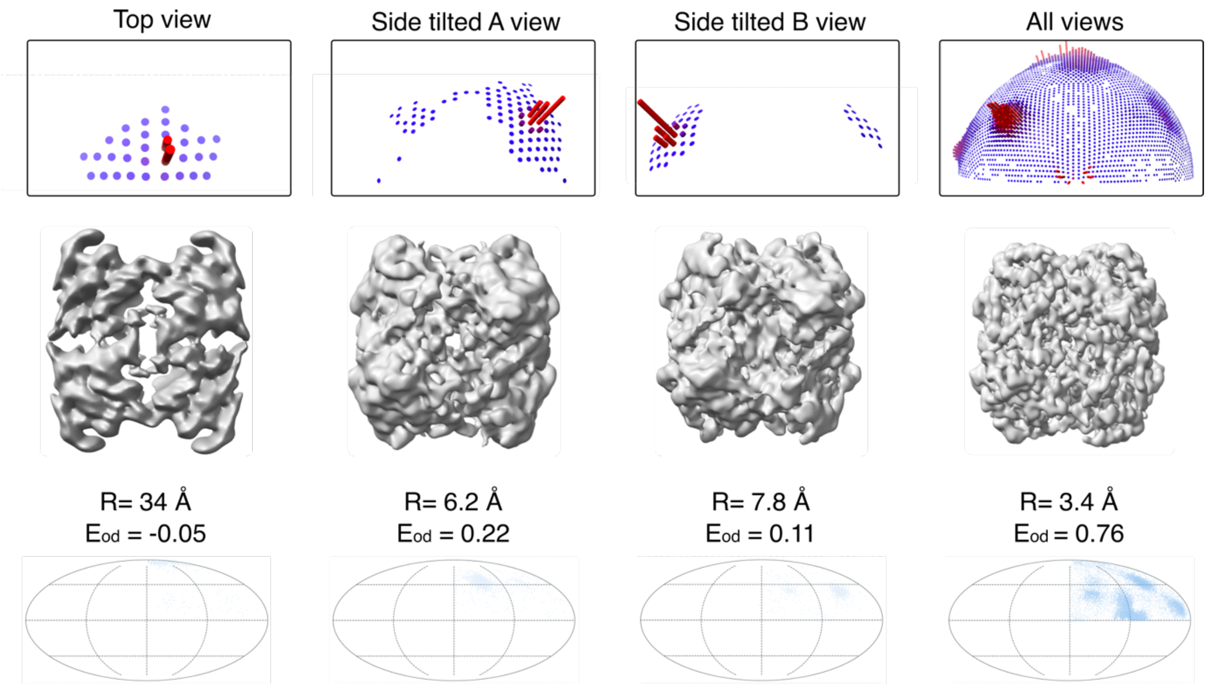


**Figure S1:** Assessment of contribution from different views to the reconstruction of catalase. Subsets of particles oriented in the same view (top, side tilted A and side tilted B) were selected from the 2D class averages from the catalase 20°C dataset without additive (Figure 5A). These subsets of particles were then used to reconstruct individual maps using the same initial model as the reference. The subset with top views alone led to a worse map with a negative value of $E_{od}$ compared to the side-tilted views. The number of particles per subset is 20,000. R = Resolution of reconstruction. $E_{od}$ = efficiency of coverage of Fourier space. The lower panel shows the extent of coverage of orientations in the Fourier space for the given subset.

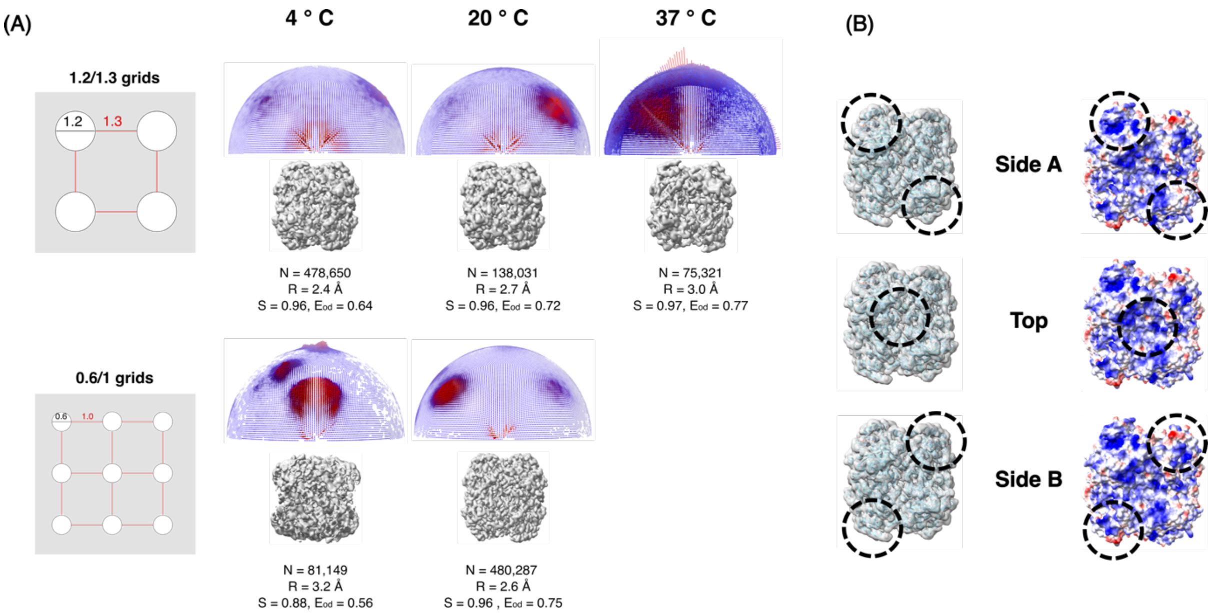


**Figure S2**: Effect of changes in orientation distribution of catalase on different grid types. (A) The differences in hole size and spacing are shown in a schematic representation of a grid square for the two types of grids used. On the right are the orientation distribution plots from Relion and cryo-EM maps of catalase frozen with different types of grids and at different temperatures. (B) Different views of catalase are marked in the figure on the left, where the dotted circle represents the location of the area of interest on the catalase map. The right panel shows the electrostatic surface representation of the map with the preferred views marked. Different parameters used to analyse the quality of the maps are shown next to the orientation plots. N = no. of particles used for reconstruction, R = resolution, S = sphericity, $E_{od}$ = efficiency.

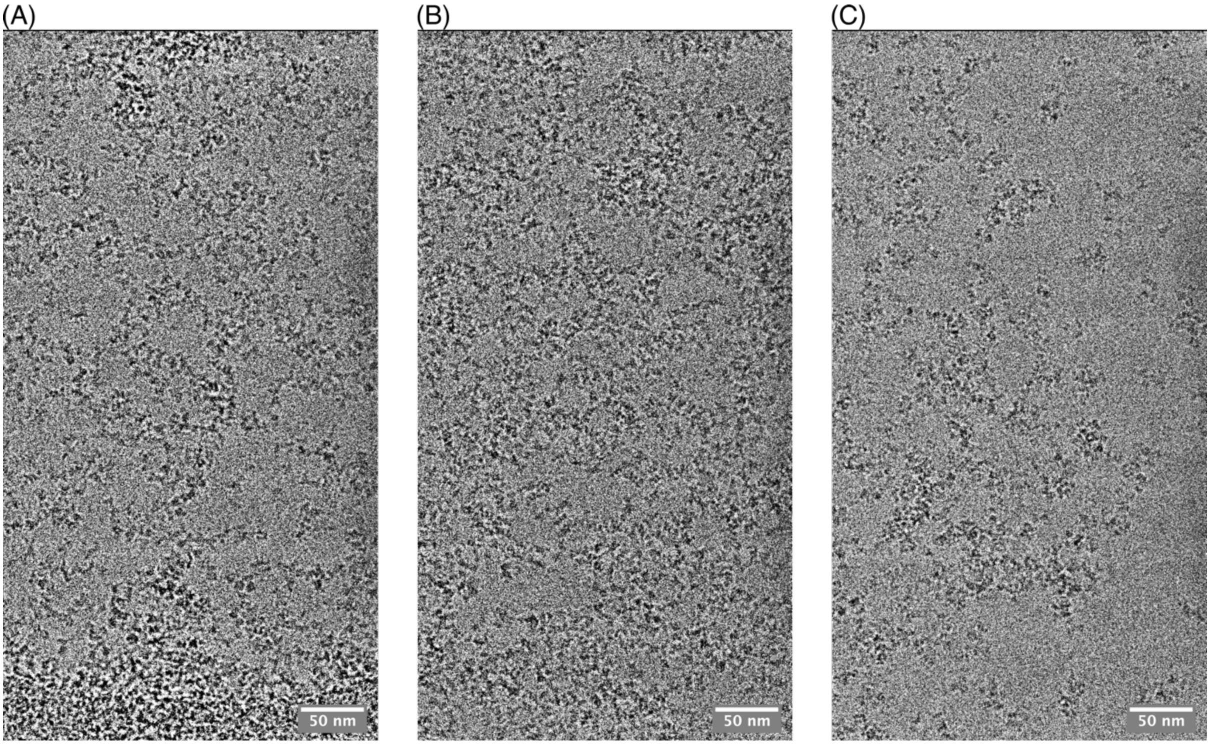


**Figure S3:** Effect of surfactant addition to PaaZ grids prepared at 20°C. (A) In the absence of any additive, PaaZ on grids shows a preference for the side view and occasional clustering of two or more hexameric PaaZ molecules (B) Upon the addition of SLS, more clustering or clumping of the PaaZ molecules was observed when compared to the no-additive condition. The hexameric PaaZ molecules are preferentially oriented similar to no additive but the severity seems to increase. (C) Upon addition of tween 20 to the sample buffer, the micrographs show clumping and aggregation of PaaZ molecules on grids.

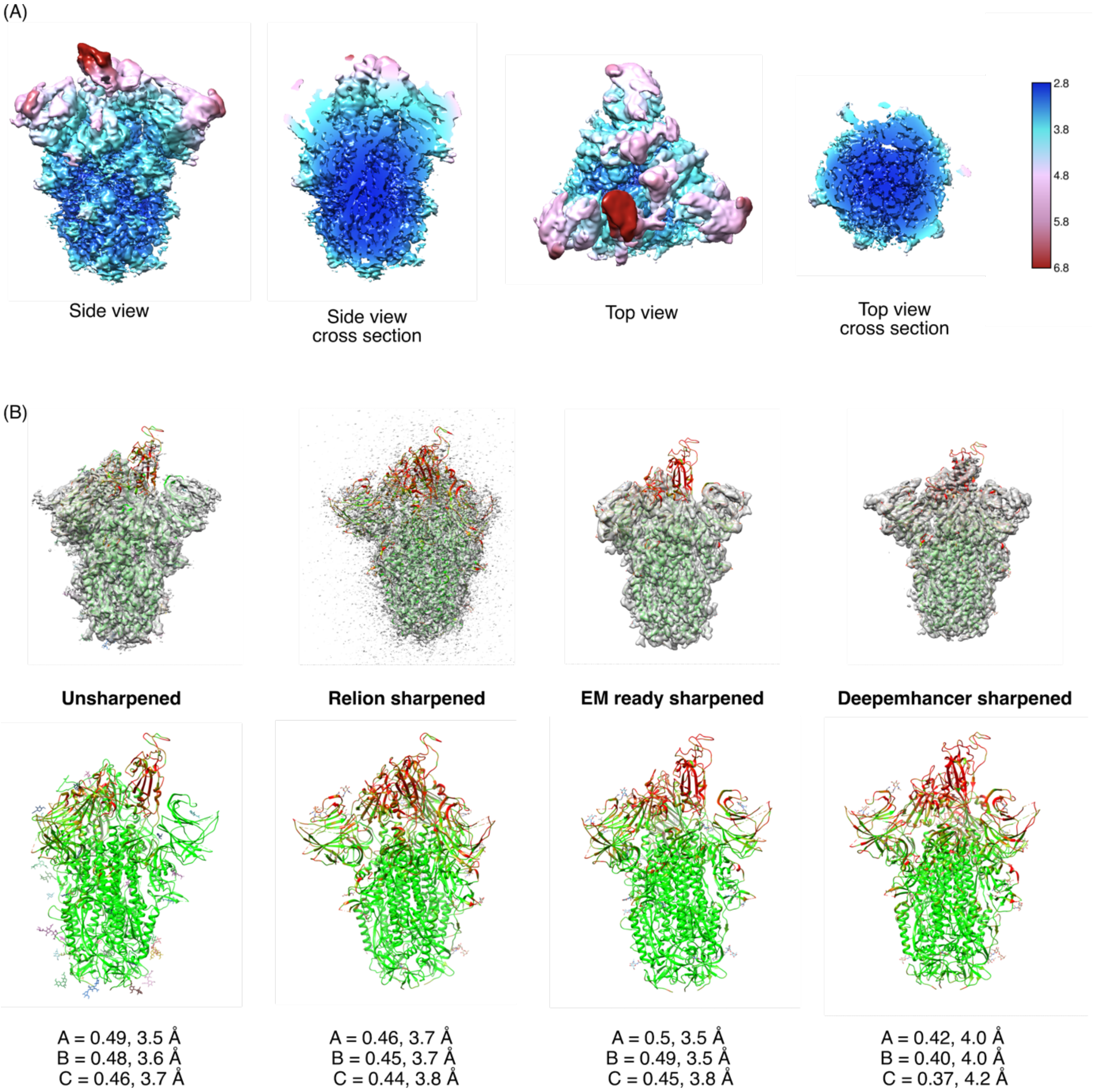


**Figure S4.** (A) Local resolution plot of map sharpened with a B-factor of -50 Å$^2$. Different views of the map show the resolution spread, which is higher at the core and lower at the periphery. The different views and cross-sections show the three RBDs at differential resolution. The colour key is shown on the right where the resolution is mentioned in Å. (B) A gallery of maps generated with different tools is coloured in transparent grey and the fit model is depicted in cartoon representation and coloured according to the Q score (Pintille et al 2020) is shown in the top panel. The threshold for the maps used are unsharpened – 0.0171, Relion sharpened with B=-50 Å$^2$ – 0.048, EMready– 2.23, and DeepEMhancer – 0.0302. In the bottom panel, the model alone is shown in cartoon representation and is coloured according to the Q score using Chimera. The Q-scores and estimated resolution by MapQ in Å for individual chains in each map are also given. The Q-score differs by the type of the maps used for calculation but uniformly, they show that the RBD is of lower resolution, also reflected in the local resolution plot shown in panel A.

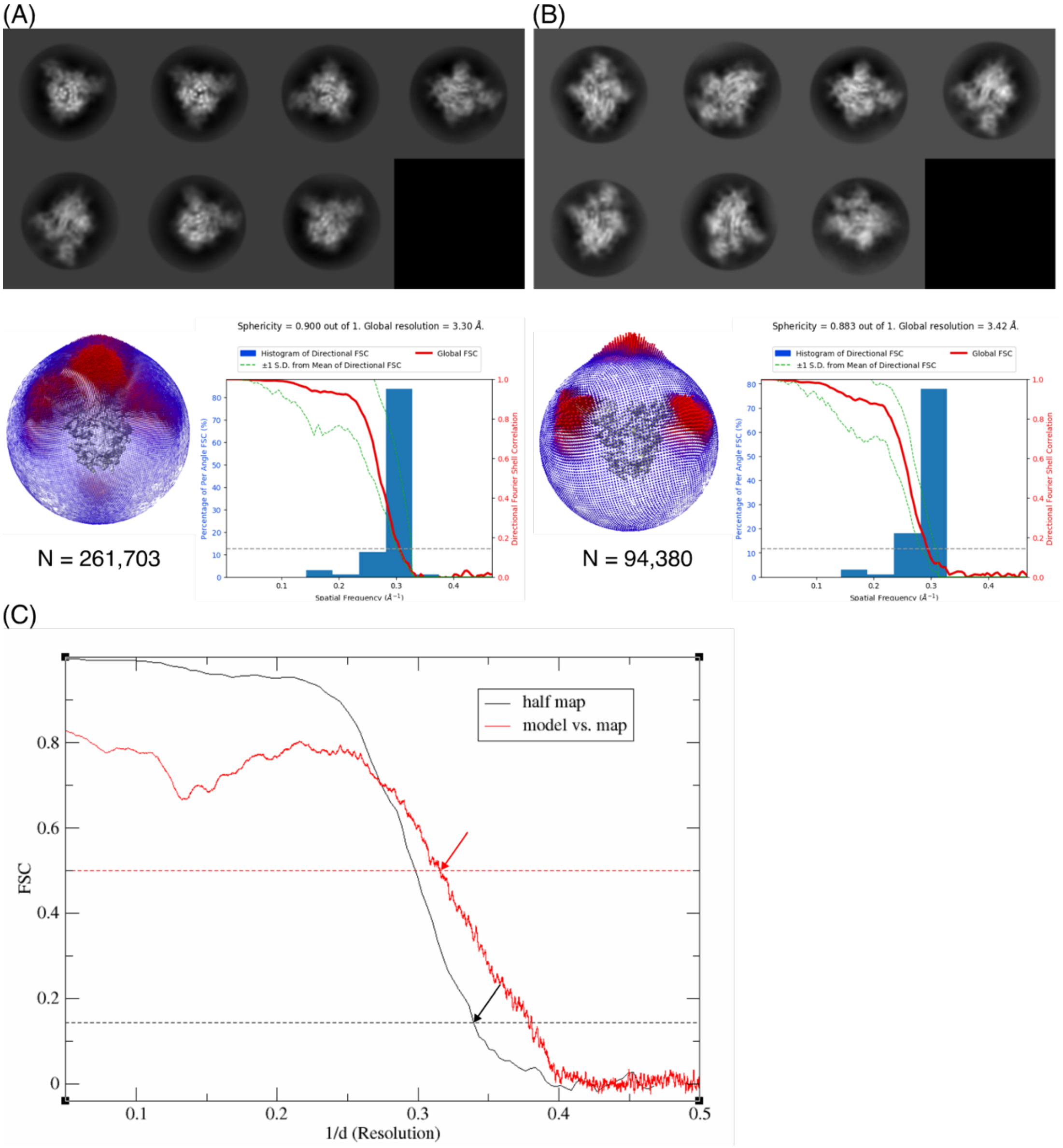


**Figure S5:** Assessment of contribution from different views to the reconstruction of the SARS-CoV-2 spike protein. (A) Representative 2D class averages, orientation distribution and 3D FSC plots of the SARS-CoV-2 spike protein without tag and with CTAB as an additive. The full dataset with all good classes was used for reconstruction, including the highly abundant top/bottom views. (B) Representative 2D class averages, orientation distribution and 3D FSC plots when only tilted-side views were used for reconstruction. These comparisons are performed before the CTF refinement and polishing on the particles. N = number of particles used for refinement and reconstruction. (C) Comparison of half-map FSC with the map model FSC of the final reconstructed map. The estimated resolution from the half map FSC (@0.143) is 3 Å and the estimated resolution obtained from the map vs. model FSC (@0.5) is 3.4 Å. The 0.143 and 0.5 FSC cut-offs are indicated by black and red lines respectively.

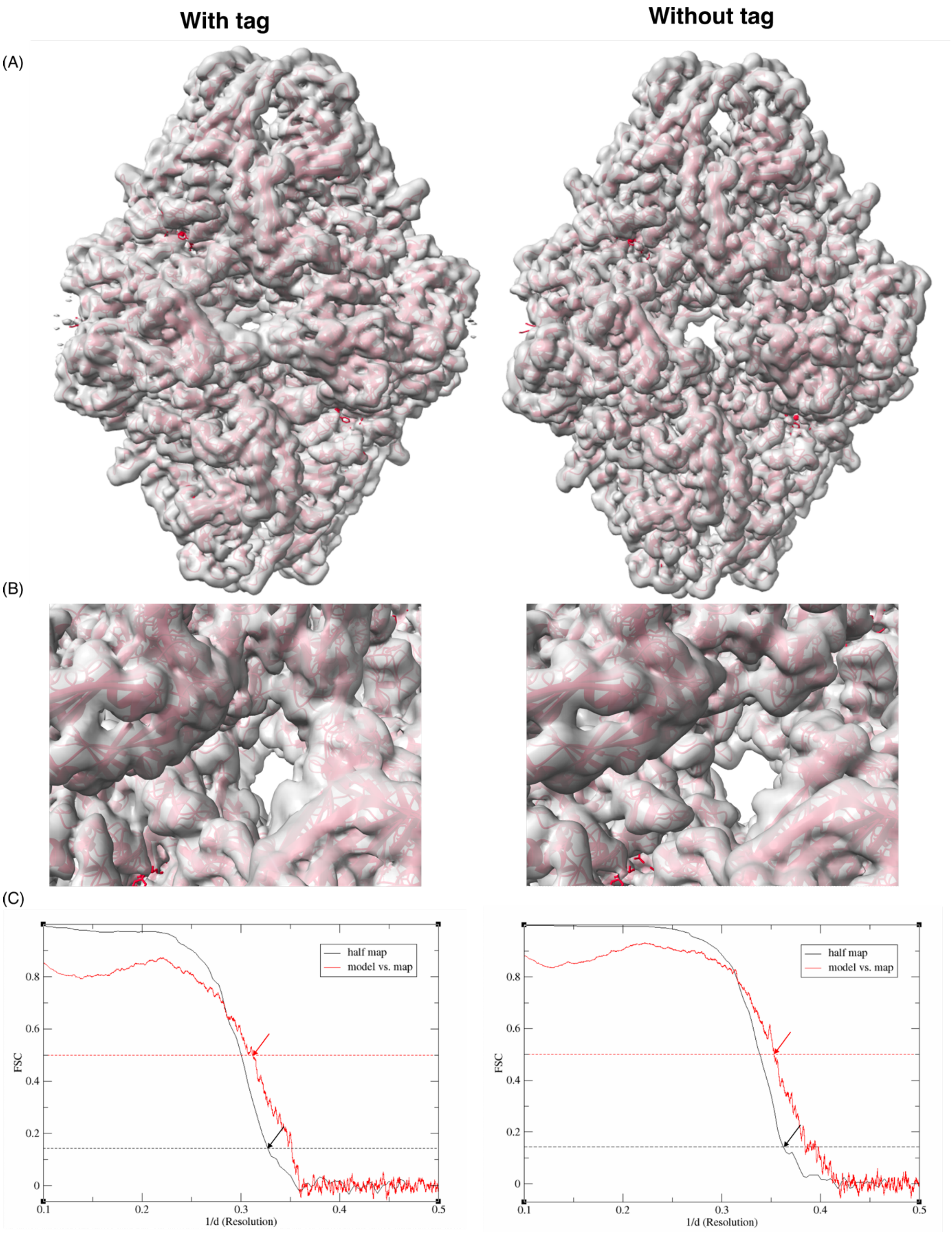


**Figure S6.** β-galactosidase map quality comparison of reconstructions with and without his-tag. (A) The unsharpened combined final maps of β-gal reconstructed from the dataset with the enzyme tagged and untagged respectively show differences in the quality of the maps. The threshold used to visualise the maps is 0.0107. (B) Zoomed-in areas of the respective maps with the model fitted (PDB ID: 6cvm) highlight the differences in quality of the maps (C) The estimated resolutions from the half map FSC

(@0.143, black line) are 3.1 Å and 2.8 Å and the map vs. model FSC (@0.5, red line) are 3.2 Å and 2.8 Å for β-gal with and without tag respectively.

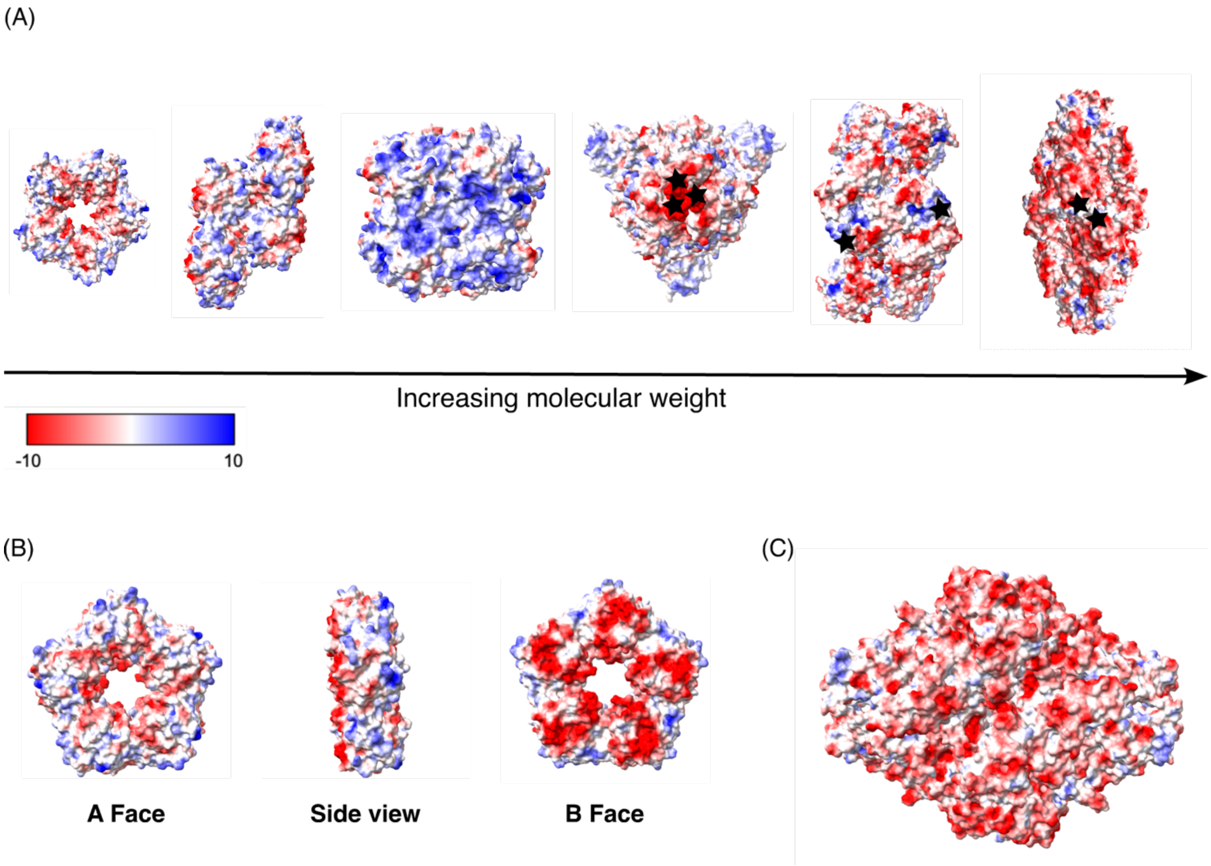


**Figure S7**: Electrostatic surface representation of the different proteins used in this study. Panel A shows human CRP-pentamer, human CRP-decamer, human erythrocyte catalase, SARS-CoV-2 spike protein, PaaZ and β-galactosidase (from left to right). Maps were generated using ChimeraX and default thresholds of -10 to +10 kcal/mol.e. were used for the colouring. The black stars in the spike protein, PaaZ and β-galactosidase indicate the location of the poly-histidine affinity tag on the respective structures. (B) Electrostatic surface representation of different views of CRP pentamer (C) Electrostatic surface representation of the β-galactosidase enzyme alternate surface.

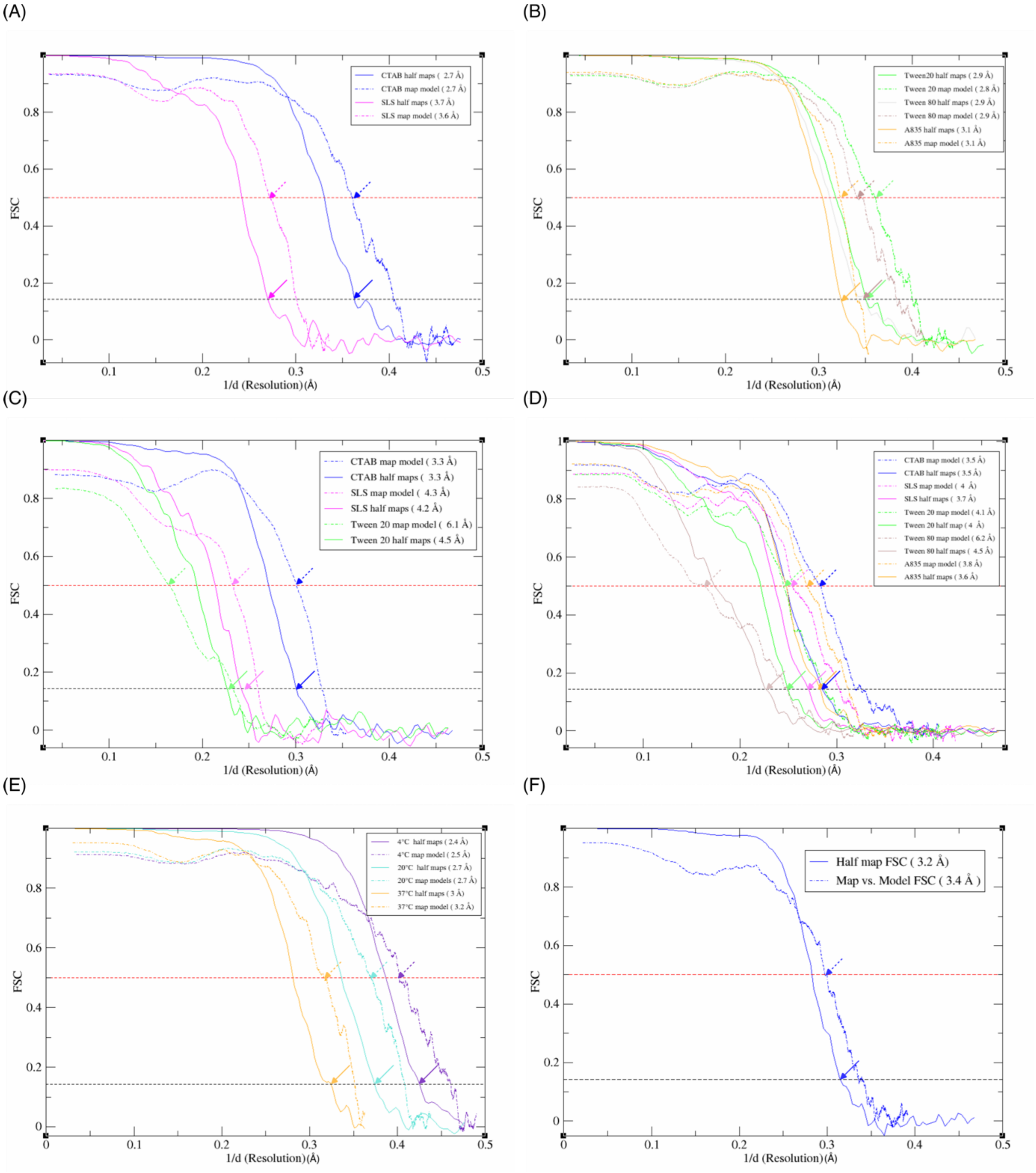


**Figure S8:** Estimation of the resolution of the cryo-EM reconstructions with half-maps and the full map against the model. (A) Catalase datasets with charged surfactant additives; (B) Catalase datasets with neutral surfactant additives; (C) CRP-pentamer datasets with surfactant additives; (D) CRP-decamer datasets with surfactant additives; (E) Catalase datasets from grids prepared at different temperatures; (F) PaaZ grids prepared at 20°C in the absence of an additive. The FSC cut-off at which the resolutions are estimated (@0.5 and @0.143 thresholds for map vs model and half-maps respectively) is marked with arrows. The values of the resolution estimates are summarised in Table S2.

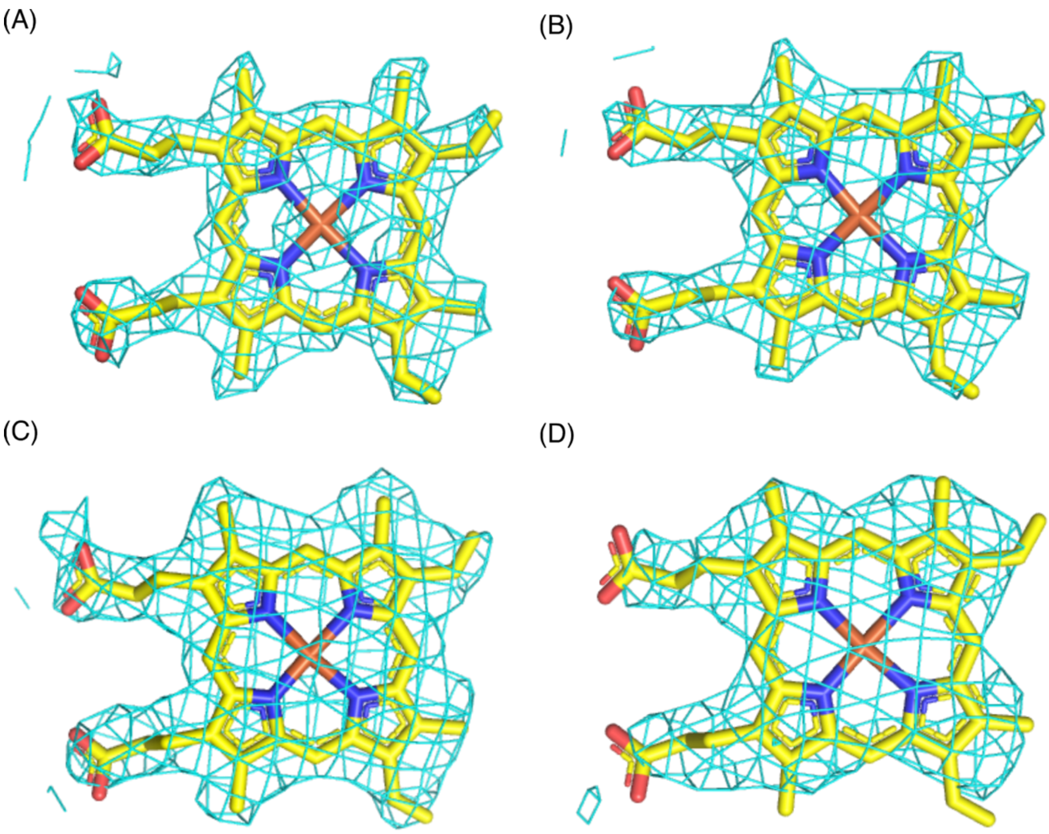


**Figure S9**: Example density from catalase datasets around the heme group. The carbon atoms of the ligand heme are in yellow and the EM map is in aquamarine. (A) 4 °C (B) 20 °C (C) CTAB (D) SLS. Pymol was used to make the final figure using the refined models and sharpened maps by carving 2. 0 Å around the atoms at 7.0 σ.

**Table S1.** The relative abundance of CRP pentamer and decamer populations in cryo-EM datasets with different additives. The numbers in the table are after two rounds of 2D classification.

| Sl. No. | Condition | % Decamer | No. of Decamer | % Pentamer | No. of Pentamer | Protein concentration |
|---|---|---|---|---|---|---|
| 1 | No additive | 16% | 9947 | 84% | 52536 | 2.1 mg/ml |
| 2 | CTAB | 42% | 25,992 | 58 % | 36,353 | 4.1 mg/ml |
| 2 | Tween 20 | 41.5% | 51784 | 58.5% | 72376 | 3.6 mg/ml |
| 3 | SLS | 52.2% | 59211 | 47.8% | 54191 | 2.6 mg/ml |
| 4 | A8-35 | 55.7% | 99908 | 44.3% | 79342 | 3.6 mg/ml |
| 5 | Tween 80 | 46.15% | 37327 | 53.85% | 43550 | 6.8 mg/ml |

**Table S2.** Summary of resolution estimates from half-map FSCs and map vs. model FSCs of protein imaged and processed in different conditions

| Protein | Condition | Resolution estimate (Å) FSC half map (@ 0.143) | Resolution estimate (Å) FSC map vs model (@ 0.5) |
|---|---|---|---|
| CRP pentamer | CTAB | 3.3 | 3.3 |
| | SLS | 4.2 | 4.3 |
| | Tween 20 | 4.5 | 6.1 |
| CRP decamer | CTAB | 3.5 | 3.5 |
| | SLS | 3.7 | 4 |
| | Tween 20 | 4 | 4.1 |
| | Tween 80 | 4.5 | 6.2 |
| | A8-35 | 3.5 | 3.7 |
| Catalase | CTAB | 2.7 | 2.7 |
| | SLS | 3.7 | 3.6 |
| | Tween 20 | 2.8 | 2.9 |
| | Tween 80 | 2.9 | 2.9 |
| | A8-35 | 3.1 | 3.1 |
| | 4°C, 1.2/1.3 grids | 2.4 | 2.5 |
| | 20°C, 1.2/ 1.3 grids | 2.7 | 2.7 |
| | 37°C, 1.2/1.3 grids | 3.0 | 3.2 |
| PaaZ | 20°C | 3.2 | 3.4 |
| | 4°C, CTAB | 2.3 | 2.4 |
| Spike | Without tag, CTAB | 3 | 3.4 |
| β-galactosidase | With tag | 3.1 | 3.2 |
| | Without tag | 2.8 | 2.8 |
